# Cardiomyocyte pyruvate dehydrogenase kinase 1 knockout activates Rho-mediated remodeling and decreases fatty acyl availability in female mice

**DOI:** 10.64898/2026.07.30.741875

**Authors:** Michael G. Atser, Sareh Panahi, Justin J. Hong, Xiaoke Hu, Aura Balita, Haoning Howard Cen, Sing-Young Chen, Vincent R. Richard, Timon Geib, Armando Alcazar Magana, Brian Rodrigues, Leonard J. Foster, Elizabeth J. Rideout, Christoph H. Borchers, James D. Johnson

## Abstract

Pyruvate dehydrogenase kinase 1 (PDK1) inhibits pyruvate dehydrogenase (PDH) and therefore may regulate the balance between glucose and fatty acid oxidation in cardiomyocytes. We assessed metabolism and function in tamoxifen-inducible, cardiomyocyte-specific *Pdk1* knockout (*Pdk1cKO*) mice, including both males and females fed a high fat diet, to resolve the role of PDK1 in the heart. Female, but not male, *Pdk1cKO* hearts showed the expected increase in cardiac PDH activity. Though cardiac function was largely preserved, female *Pdk1cKO* hearts had smaller left ventricles. Proteomics and phosphoproteomics revealed no change in classical hypertrophic markers but downregulated aerobic respiration alongside upregulated Ca^2+^ handling and Rho signalling, suggesting metabolic remodeling with altered cardiomyocyte contractility and cytoarchitecture in *Pdk1cKO* hearts. Concordantly, lipidomics revealed membrane remodeling marked by elevated saturated phosphatidylcholine in female *Pdk1cKO* hearts and decreased triacylglycerols, free fatty acids and fatty acid uptake proteins (CD36, FABP3). These findings reveal novel, female-predominant cardiometabolic roles for PDK1.

## Introduction

Metabolism is essential to cardiac function and is a critical factor that distinguishes a healthy heart from a diseased one. The heart can use various nutrient substrates for energy, but primarily uses glucose and fatty acids under physiological conditions^1^. Flexibility between glucose and fatty acid oxidation is critical for cardiac physiology and depends on energy demand, hormonal status, nutrient availability, and oxygen availability. The functional consequences of metabolic flexibility depend on metabolic context. For example, reduced oxygen availability in the ischemic heart causes an initial increased reliance on glucose metabolism, mediated through increased glucose uptake^2^. Chronic ischemia results in a further shift to anerobic glucose metabolism^3^, mediated by pyruvate dehydrogenase (PDH) suppression^4^, increasing acidosis and exacerbates cardiac dysfunction^5^.

The consequences of cardiac dysmetabolism are sexually dimorphic, in line with the differences in incidence and progression of cardiometabolic disease in women and men^6^. Genetically engineered mouse model generally show increased sensitivity to cardiometabolic disease in males compared to females^7^. Examples of male-biased effects include: 1) deletion of *Ppara*, a master lipid regulator, driving cardiac triacylglycerol accumulation and death^8^; 2) deletion *Fkbp1b*, responsible for cardiomyocyte Ca^2+^ handling, on cardiac hypertrophy^9^; 3) deletion of myosin heavy chain on left ventricular dysfunction^10^. Nonetheless, cardiometabolic disease has a higher prevalence in females^7,11^. Despite these clear and impactful differences, the effects of biological sex on cardiomyocyte metabolism remain understudied.

The pyruvate dehydrogenase kinase (PDK) family contains four isoforms that regulate cardiomyocyte substrate preference. Canonically, they inhibit glucose oxidation by phosphorylating and inactivating pyruvate dehydrogenase (PDH) resulting in the inhibition of pyruvate decarboxylation to acetyl-CoA for entry into the citric acid cycle. The PDK1 isoform is abundant in the heart and has been implicated in cardiovascular disease and cardiac metabolism. According to the GTEX database, *PDK1* mRNA expression is highest in heart relative to all other adult tissues. Genetic variation at the *PDK1* locus is moderately associated with heart failure, with a greater association than the other PDK isoforms, according to the Common Metabolic Disease Knowledge Portal (CMDKP; hugemap.org). *PDK1* mRNA expression is upregulated in atherosclerosis and arrhythmogenic right ventricular cardiomyopathy^12–14^. The PDK1-specific ser232 site on PDH was reported to be dephosphorylated in diabetic and ischemic cardiomyopathic patients^15^. However, we recently reported the surprising result that PDK1 did not affect PDH activity in H9c2 cells but instead supported triacylglycerol hydrolysis in H9c2 cells^16^. Consistent with a novel, non-canonical role in lipid metabolism, lifelong whole-body *Pdk1* knockout resulted in impaired triacylglycerol hydrolysis and cardiac triacylglycerol accumulation in male mice^16^. However, the *in vivo* metabolic phenotype of acute, cardiomyocyte-specific PDK1 knockout is unclear in both males and females. In this study, we used an inducible, tissue-specific knockout model to assess the non-canonical roles of PDK1 on cardiac function using echocardiography and metabolism using proteomics, phospho-proteomics, and lipidomics/metabolomics in male and female mice.

## Results

### Gene dosage-dependent reduction of Pdk1 mRNA and protein in male and female Pdk1cKO hearts

To study the specific role of PDK1 in heart function and metabolism, we knocked out the *Pdk1* gene in mouse cardiomyocytes (*Pdk1cKO*) by breeding heterozygous floxed mice containing loxP sequences flanking the second exon of the gene^17^ with tamoxifen-inducible Cre mice under the influence of the cardiomyocyte-specific *Myh6* promoter to generate wildtype Cre controls (*Pdk1*^wt/wt^;*Myh6*Cre^ERT^), *Pdk1*cHET (*Pdk1*^fl/wt^;*Myh6*Cre^ERT^), and *Pdk1cKO* (*Pdk1*^fl/fl^;*Myh6*Cre^ERT^) mice. The offspring were weaned onto a high fat diet at 3 weeks and then injected with either corn oil as a vehicle control or tamoxifen to induce activation of Cre recombinase at 8-10 weeks (Fig. 1A). Two weeks after tamoxifen injection, there was a decrease in *Pdk1* mRNA expression in *Pdk1*cKO hearts relative to control hearts of both male and female mice (Fig. 1B). We observed no changes in mRNA expression of the *Pdk*2, 3 and 4 isoforms (Fig. 1C-E). We used tamoxifen-injected Cre-expression mice injected as controls (CreTam) to control for any effects of Cre^18^ or tamoxifen^19^. They showed no difference in *Pdk* mRNA expression relative to corn-oil-injected Cre-expression mice (Fig. S1A-D). Mass spectrometry revealed PDK1 protein abundance was significantly reduced in whole male (-67%) and female (-75%) *Pdk1*cKO hearts (Fig. 1F). Residual PDK1 protein detection was expected because whole hearts contain other cell types that would be experience cardiomyocyte-specific Cre activity. Other PDK isozymes showed no changes in protein abundance except for PDK2, which showed a 30% decrease in female *Pdk1*cKO mice hearts (Fig. 1G-I). There were no changes PDK isoform mRNA levels in liver or gastrocnemius skeletal muscle, metabolically relevant tissues with significant *Pdk1* expression^20^(Fig. S1E-L). These observations show cardiac-specific *Pdk1* knockout occurs within 2 weeks of tamoxifen administration.

**Figure 1.**
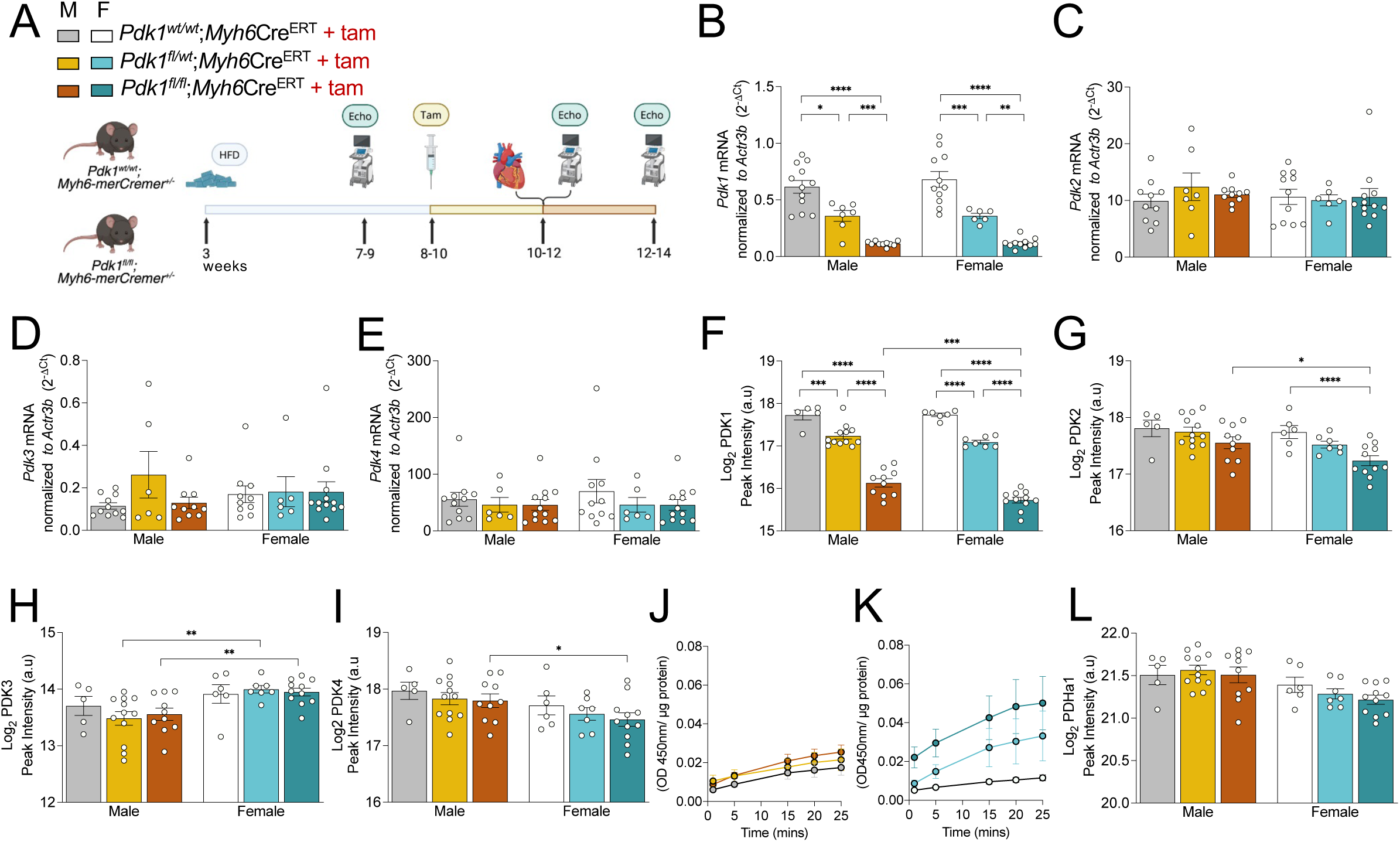
Validation of cardiomyocyte *Pdk1* knockout model. **(A)** Schematic of experimental design and legend for subsequent panels. **(B-E)** *Pdk1*, *Pdk2*, *Pdk3*, and *Pdk4* mRNA expression in whole heart, 2 weeks after tamoxifen injection. **(F-I)** PDK1, PDK2, PDK3, PDK4 protein abundance measured by LC- MS/MS proteomics in whole hearts, 2 weeks after tamoxifen injection. **(J)** PDH activity in male and **(K)** female mice hearts. **(L)** PDHa1 protein abundance measured by LC-MS/MS proteomics. Data are shown as mean ± SEM and were analyzed by either a Two-way ANOVA followed by a post-hoc Tukey multiple comparison test (B-I; L) or linear mixed effects model where time and genotype were fixed effects and the replicates was a random effect (J-K). ^∗^ *p* < 0.05; ^∗∗^ *p* < 0.01; ^∗∗∗^ *p* < 0.001; ∗∗∗∗ *p* < 0.0001.

We measured the impact of cardiac-specific PDK1 reduction on cardiac PDH activity (Fig. 1J-K). We observed a non-significant time x genotype interaction in males (F(1,28) 3,22, *p* = 0.08), suggesting no differential activity between controls and *Pdk1*cKO. In contrast, there was a significant time x genotype interaction in females (F(1,25) = 16.1, *p* < 0.001), indicating differential PDH activity that was elevated in female *Pdk1*cKO mice compared to control (Fig. 1K). PDH protein abundance was unaltered in *Pdk1*cKO mice of both sexes, indicating that elevations in PDH activity were not a consequence of increased abundance (Fig. 1L). Together, these observations reveal a sex-specific differential response in cardiac PDH activity following acute, cardiac-specific *Pdk1* reduction, wherein females show elevated PDH activity in contrast to males, whose activity remain unaltered.

### Female hearts have reduced left ventricular volume but are otherwise normal 4 weeks after Pdk1 ablation

We took baseline echocardiography measurements ∼1 week before tamoxifen injection and subsequent measurements, 2 and 4 weeks after tamoxifen injection (Fig. 1A). Stroke volume and cardiac output were unaltered in *Pdk1*cKO mice of both sexes (Table 1). We found no changes in the early to late peak velocity (E/A) and early peak to annular velocity (E/E’) ratios in male or female *Pdk1*cKO mice following tamoxifen injection (Table 1, Fig. S2). However, female *Pdk1*cKO mice had smaller left ventricular internal diameter and volume compared to controls at the 4-week time point (Table 1, Fig. S2F-I). Although male *Pdk1*cKO mice showed similar trends as in females, they were not significantly different (Fig. S2F-I). There was no significant change in left ventricular posterior wall thickness and interventricular wall thickness in *Pdk1*cKO mice of both sexes (Table S1). Together, these data show that cardiac function in male and female mice is largely resistant to acute PDK1 reduction, although female mice exhibited consistent morphological changes.

**Table 1.** Cardiac function in acute cardiomyocyte-specific *Pdk1* knockout mice. Change from baseline at 4 weeks (4 wks – baseline). Data are shown as mean ± SEM and were analyzed by an unpaired two-tailed Student’s t-test. Female CreTam control n = 3, male CreTam control n = 4, female *Pdk1*cKO n = 6, male *Pdk1*cKO n = 3. ^∗^ *p* < 0.05.

|  | **Control** |  | ***Pdk1cKO*** |  |
| --- | --- | --- | --- | --- |
|  | Male | Female | Male | Female |
| Cardiac output | -0.41 ± 2.35 | 2.44 ± 1.33 | 4.49 ± 2.50 | -2.08 ± 1.42 |
| Stroke volume | 2.18 ± 2.95 | 4.52 ± 1.52 | 7.64 ± 6.07 | -1.01 ± 3.83 |
| Ejection fraction | -7.75 ± 2.3 | -12.64 ± 2.8 | 4.67 ± 6.5 | -5.37 ± 5.2 |
| Fractional shortening | -5.68 ± 1.9 | -8.78 ± 1.9 | 3.69 ± 4.9 | -4.07 ± 3.9 |
| Early/late peak velocity (E/A) | -0.06 ± 0.23 | 0.09 ± 0.35 | 0.25 ± 0.05 | 0.16 ± 0.32 |
| Early peak/annular velocity (E/E') | -2.53 ± 5.07 | -7.55 ± 8.83 | 7.21 ± 5.49 | 0.80 ± 6.52 |
| Diastolic left ventricular internal diameter | 0.31 ± 0.11 | 0.54 ± 0.12 | 0.27 ± 0.28 | **-0.02 ± 0.13\*** |
| Systolic left ventricular internal diameter | 0.38 ± 0.06 | 0.69 ± 0.05 | 0.01 ± 0.29 | **0.13 ± 0.17\*** |
| Diastolic left ventricular volume | 11.41 ± 3.78 | 21.43 ± 5.22 | 9.39 ± 8.89 | **-1.02 ± 4.48\*** |
| Systolic left ventricular volume | 8.07 ± 1.36 | 16.00 ± 2.19 | -0.05 ± 5.39 | **2.00 ± 3.45\*** |

### Proteomic analysis of Pdk1cKO hearts

We analyzed the cardiac proteome at 2 weeks post tamoxifen injection, as the timepoint with minimal morphological or functional differences. We fasted mice for 4 h and isolated hearts thereafter, since overall PDK activity increases during fasting^21–23^. We annotated 5640 proteins across all groups. Partial Least Squares-Discriminant Analysis (PLS-DA) using sex and genotype as response variables revealed modest separation of clusters, with some overlap (Fig. 2A). Interestingly, we observed that the male and female CreTam controls were somewhat distinct clusters. To further explore baseline sex differences in the cardiac proteome, we performed a differential expression analysis between male and female controls and identified 63 proteins with significantly different abundances (Fig. S3A). We identified enrichment in ribosomal processing pathways in female controls hearts and aerobic respiration and complement cascade pathways in male control hearts (Fig. S3B,C).

**Figure 2.**
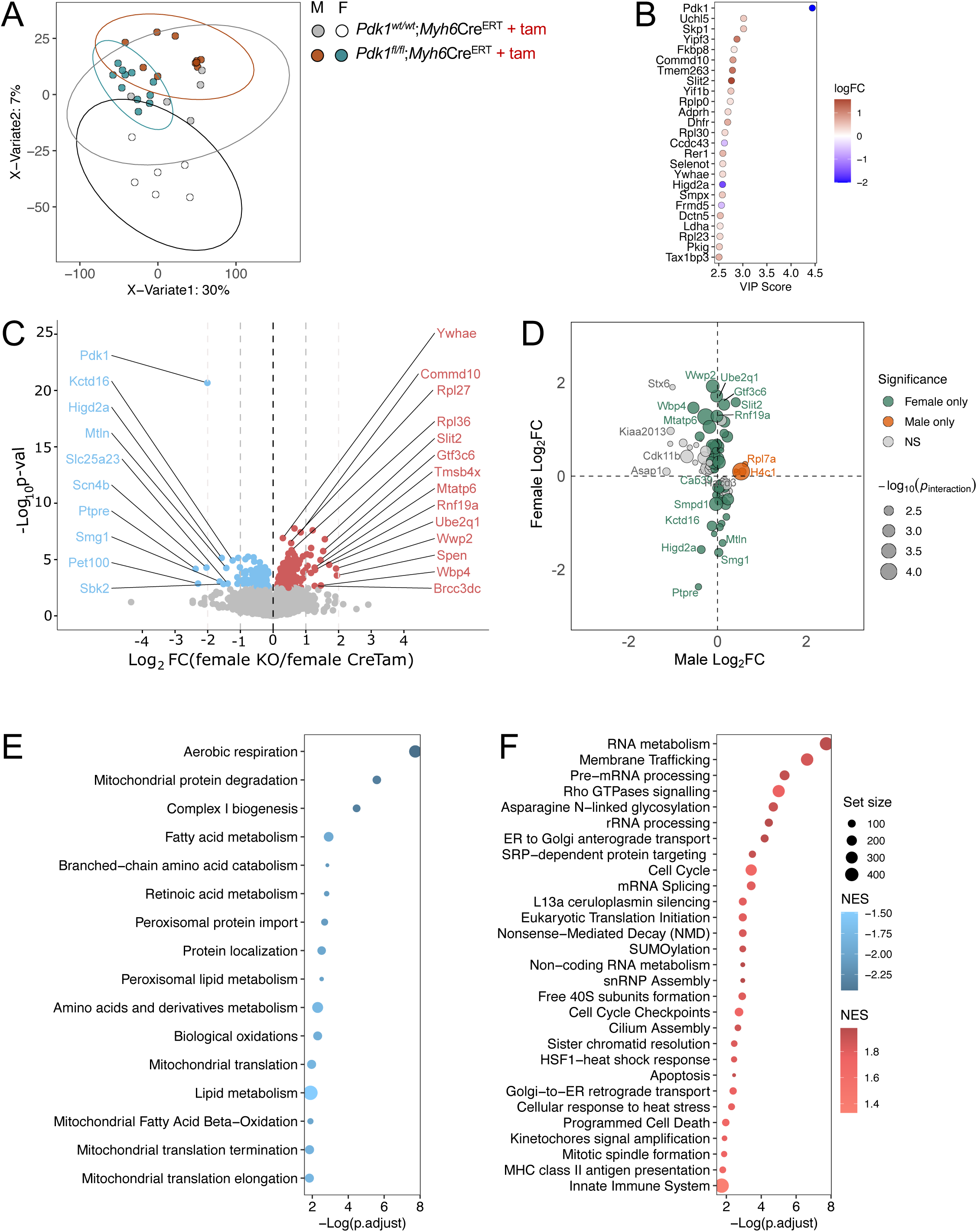
Cardiomyocyte-specific PDK1 depletion drives female-predominant remodeling of the cardiac proteome. **(A)** Partial least square discrimination analysis of protein abundances in mice hearts, 2-weeks post-tamoxifen injection. **(B)** Top 25 proteins with the highest variable importance projection component 1 scores between genotypes. Blue dots are upregulated and red dots are downregulated in female *Pdk1*cKO hearts. **(C)** Volcano plot depicting differentially abundant proteins in female *Pdk1*cKO relative to control. **(D)** Scatter plot visualising proteins with a sex-specific response to PDK1 reduction. Green dots represent proteins that were significantly changed in females. Orange dots depict proteins that were significantly changed in males. Grey dots are proteins that were not significantly altered in either male or female individual analyses. Gene set enrichment analysis based on the reactome database annotation depicting **(E)** downregulated and **(F)** upregulated pathways in female *Pdk1*cKO hearts. Data were analyzed by Limma. Female CreTam control n = 6, male CreTam control n = 5, female *Pdk1*cKO n = 11, male *Pdk1*cKO n = 10.

We next determined the proteins that were differentially abundant between *Pdk1*cKO and CreTam control in both male and female mice. Variable Importance Projection (VIP) analysis showed that PDK1 was most responsible for differences between control and *Pdk1*cKO across sexes (Fig. 2B). The second most important protein in this analysis was UCHL5, a key regulator of proteostasis that was reported to be upregulated in cardiac ischemic-reperfusion injury^24^, which was increased in female *Pdk1*cKO hearts (Fig. 2B). The third most important protein was SKP1, which is also a key regulator of protein quality control and cardiac hypertrophy^25^. Interestingly, we observed more significantly different proteins in females (359 proteins) than males (26 proteins), with 15 proteins common to both males and females (Fig. 2C, S3D,E), suggesting a more pronounced response in females. We next performed interaction analysis to determine which proteins had a sex-specific response but found no proteins with adjusted *p* < 0.05, possibly due to limited sample sizes. We resorted to a nominal *p* value threshold of < 0.01 to identify candidate proteins with a sex-specific response to PDK1 loss. We observed a clustering of proteins along the female-effect axis, with minimal changes on the male’s, indicating that proteins with strong interaction effects were predominantly driven by changes in female *Pdk1*cKO mice, with minimal genotype-associated changes in males (Fig. 2D). We determined the pathway enrichment of differentially abundant proteins in both sexes using a Gene set enrichment analysis (GSEA) method with the Reactome database annotations. We found negative enrichment in metabolic pathways such as aerobic respiration, amino acid catabolism, and lipid metabolism in females (Fig. 2E). In contrast, RNA metabolism and Rho signaling pathways were positively enriched in female *Pdk1*cKO hearts (Fig. 2F). Consistent with pathway-level enrichment, individual Rho signaling proteins showed coordinated changes in female *Pdk1*cKO hearts, marked by increased abundance of the RHOB GTPase, its activating guanine nucleotide exchange factor ARGAL, and the downstream effector ROCK1, alongside decreased abundance of the inhibitory GTPase activating protein RHG10 (Fig. S4A-D). In males, the only significantly enriched pathway was translation (upregulated). These data show that cardiac-specific *Pdk1* deletion resulted in modest but broad changes in the female *Pdk1*cKO cardiac proteomic.

To assess possible molecular mechanisms underlying the smaller left ventricular size in female *Pdk1*cKO mice we undertook a targeted mining of our proteomic data for: 1) signatures that may predispose these hearts towards hypertrophic remodeling, and 2) signatures of enhanced intrinsic contractility marked by cytosolic Ca^2+^ elevation that could maintain cardiac output and stroke volume from a smaller ventricular volume. We observed no significant changes in established cardiac hypertrophy markers ANF, MYH7 and ACTS in *Pdk1*cKO mice of both sexes (Fig. S4E-G). We also observed no significant changes in protein levels of the voltage dependent L-type Ca^2+^ channels or Ryr2 Ca^2+^ channels (Fig. S4H,I). Female *Pdk1*cKO hearts did show increased abundance in the Ca^2+^ sensor calmodulin (CALM) as well as a tendency for higher abundance of troponin C (TNNC1; *p* = 0.052)(Fig. S3J,K). Female *Pdk1*cKO hearts also showed elevated abundance of the Na^+^/Ca^2+^ exchanger NAC1 and a downregulation of phospholamban (PLN) (Fig. S4L,M), which inhibits the SERCA pump that requesters Ca^2+^ into the sarcoplasmic reticulum. These signatures suggest enhanced Ca^2+^ buffering capacity and clearance in female *Pdk1*cKO hearts. It is possible that maintenance of cardiac output may rely more on increased myofilament Ca^2+^ sensitivity and elevated Ca^2+^ cycling than on elevated peak cytosolic Ca^2+^.

### Phosphoproteomic analysis of Pdk1cKO hearts

Since PDK1 has been primarily known for its role as a kinase, we determined the effect of cardiac-specific ablation of the kinase on the phosphoproteome. We annotated 4543 phosphosites corresponding to 2186 phosphoproteins and normalized phosphosite abundance by their corresponding protein abundance. A PLS-DA visualizing the multivariate relationship between sex and genotype showed substantial overlap of male and female CreTam control clusters (Fig. 3A), indicating broadly similar phosphorylation profiles. Clusters showed larger separation, albeit still overlapping, in *Pdk1*cKO males and females (Fig. 3A), indicating that PDK1 reduction introduces some sex-dependent differences in the phosphorylation profile. The phosphosite most responsible for the difference between groups (highest VIP score; Fig. 3B) was Ser2663 on A-kinase anchor protein 13 (AKAP13), which links ADRA1B adrenergic receptors to RHOA activation^26^. Also of interest was Ppp1r12b_T649 (MYPT2) and Arhgef2_S151 (ARHG2), both hyper-phosphorylated in female *Pdk1*cKO hearts (Fig. 3B,C). ARHG2 is a guanine nucleotide exchange factor isoform that induces Rho signaling through activation of the RhoA GTPase^27,28^. Phosphorylation of Ser151 increases its catalytic activity, activating Rho signaling^29^. Interestingly, activated RhoA signaling results in an inhibitory phosphorylation of MYPT2 via ROCK1 at Thr646 (Thr645 in mice) to modulate cardiac contraction^30,31^. Although Thr645 was not detected in our dataset, the neighbouring Thr649 lies within the same well-characterized ROCK-inhibitory site but has not been functionally characterized. Consistent with the phosphoproteomic evidence that MYPT2 activity may be decreased in female *Pdk1*cKO hearts, we observed proteomics evidence of a reduction in MYPT2 (Ppp1r12rb) protein abundance (Fig. S4N). Together with the clear effect on AKAP13_S2663, we interpreted these data to be consistent with activated Rho signaling in female *Pdk1*cKO hearts. Overall differential phosphorylation analysis showed that female hearts had 283 differentially phosphorylated sites, while male hearts had only 36 (Fig. 3C, S5A-B). Tpm1_Y162 and Eif4g3_T253 stood out in this analysis as dephosphorylated sites with the strongest confidence. Though functionally uncharacterized, EIF4G3 is involved in mRNA cap recognition, critical for translation initiation^32^ and linked to congenital heart disease when silenced^33^. On the other hand, TPM1 (tropomyosin alpha-1 chain) stabilizes actin filaments to regulate sarcomere contraction^34^. Missense mutations in the *TPM1* gene can cause familial cardiomyopathy^35^ characterized by deterioration of Ca^2+^ transients^36^.

**Figure 3.**
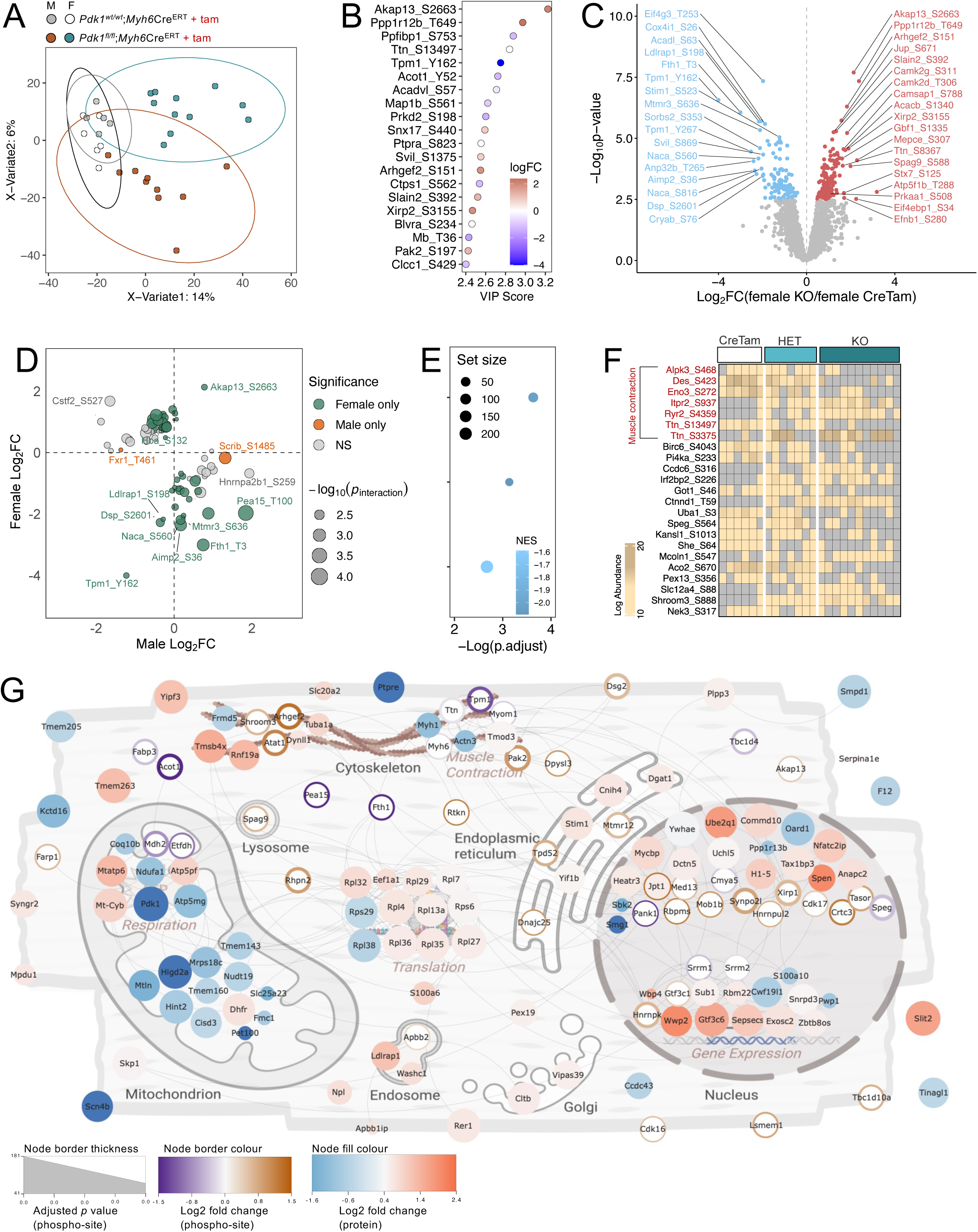
PDK1 deficiency causes sex-specific cardiac phosphoproteomic changes in mice. **(A)** Partial least square discrimination analysis of protein-normalized phosphosite abundances in mice hearts, 2-weeks post-tamoxifen injection. **(B)** Top 20 phosphosites with the highest variable importance projection component 1 scores. Blue dots are dephosphorylated, red dots are hyperphosphorylated, and white dots are not significantly altered in female *Pdk1cKO* hearts. **(C)** Volcano plot depicting differentially abundant phosphosites in female knockouts relative to control. **(D)** Scatter plot visualising phosphosites with a sex-specific response to PDK1 reduction. Green dots represent phosphosites that were significantly changed in females. Orange dots depict phosphosites that were significantly changed in males. Grey dots are phosphosites that were not significantly altered in either male or female individual analyses. **(E)** Gene set enrichment analysis on collapsed phosphoproteins based on the reactome database annotation depicting dephosphorylated pathways in female *Pdk1cKO* hearts. **(F)** Heatmap of phosphosites with significant missingness in female mice evaluated by Barnard’s unconditional exact test (nominal *p* < 0.05). **(G)** Subcellular mapping of statistically significant differentially regulated proteins and phosphoproteins in females. Protein-protein interactions were mapped out in STRING using a medium functional score of 0.4. Data were analyzed by Limma. Female CreTam control n = 6, male CreTam control n = 5, female *Pdk1*cKO n = 11, male *Pdk1*cKO n = 10.

We next identified the phosphosites that showed a sex-specific response following PDK1 reduction. Using a threshold nominal *p* < 0.01, we identified 86 phosphosites mostly occupying the quadrants indicative of discordant regulation between males and females (Fig. 3D). The majority of these phosphosites were only significantly changed in females (Fig. 3D). We then sought to determine the overall phosphorylation status of biological pathways. Unlike proteomics, phosphoproteomics has the unique challenge that multiple phosphosites can be matched to a single phosphoprotein. Previous methods have used different criteria in selecting a single site for the pathway enrichment. Since the majority of identified phosphosites across phosphoproteomics datasets remain uncharacterized, we opted for a different method that accounts for all identified phosphosites. We first determined the relative overall phosphorylation status of each phosphoprotein using a combination of linear models (when a phosphoprotein has a single phosphosite) and linear mixed effects model (in cases of multiple phosphosites), where protein abundance was used as covariate and phosphosites were assigned as random effects. Next, we performed a GSEA by ranking the phosphoproteins by *p* value and sign of Log_2_ fold change for directionality, with annotation based on the Reactome database. We observed a decrease in phosphorylation states of phosphoproteins associated with aerobic metabolism and electron transport (Fig. 3E). Importantly, this was not indicative of the activation state of these pathways but rather the overall phosphorylation state. No pathways were significantly (*p* adjust < 0.05) enriched in males. We ran parallel analyses with non-protein-normalized phosphoproteomics data and observed similar findings (Fig. S5C-G). We analysed the significant phosphosites we observed against their proteomic abundances and observed that generally phosphosite differences were not due to underlying protein abundance differences (Fig. S5H,I).

Our dataset had missing values in a genotype-specific pattern (Fig. 3F, S5J), suggesting the possibility of complete loss or gain of phosphosites in the context of *Pdk1*cKO. We performed an exploratory missingness test to determine if phosphosites were differentially detected between genotypes using a Barnard’s unconditional exact test. A limitation of this method is that it typically requires n’s greater than 20 to identify hits with adjusted *p* < 0.05. So, we used a threshold nominal *p* < 0.05, with at least a 70% missingness (absence) in either control or *Pdk1*cKO group. We observed that sites on phosphoproteins involved in muscle contraction showed a significant missingness in females (Fig. 3F), consistent with our quantitative findings of altered Rho signaling and Ca^2+^ handling protein abundance. Rather than reflecting graded changes in phosphorylation stoichiometry, this pattern suggests that a subset of contraction associated phosphosites may undergo virtually on/off switching between phosphorylation states in *Pdk1*cKO hearts.

To provide sub-cellular and relational context to our data, we mapped a protein-protein interaction network of the top 100 ranked differentially abundant proteins in females and differentially phosphorylated phosphoproteins to their subcellular location (Fig. 3G) using the Search Tool for the Retrieval of Interacting Genes/Proteins (STRING) database. The top differences were distributed across multiple subcellular compartments, but a majority were localized to the nucleus and mitochondria. Interestingly, most of the hits localized in the mitochondria were downregulated on either proteomic or phosphoproteomics level, in contrast to the nucleus, where most were upregulated (Fig. 3G). These phosphorylation changes are unlikely to be directly mediated by PDK1 kinase activity, given that all evidence points to extreme substrate specificity (PDHA1/2). Indirect mechanisms therefore likely include signalling through altered levels of lipids or metabolites.

### Lipidomic and metabolomic analysis of Pdk1cKO hearts

We have reported that PDK1 dysregulation disrupts triacylglycerol hydrolysis and the broader lipidome in a cardiomyocyte-surrogate cell line^16^. To determine the impact of PDK1 loss on the cardiac lipidome *in vivo*, we annotated 555 lipid features spanning 17 lipid classes from MS/MS spectra (Fig. 4A). We included both tamoxifen control (lacking Cre) and Cre control (injected with tamoxifen). Permutational multivariate analysis of variance (PERMANOVA) confirmed that lipidomes were not significantly different between both female control groups (tamoxifen-injected control lacking Cre; tamoxifen and Cre control; R^2^ = 0.04111; *p* = 0.725). Similarly, only 8 lipids were different between these controls using limma analysis (Fig. S5A). In males, there were significant differences between controls (R^2^ = 0.38549; *p* = 0.043). Fifty-two lipid species were identified to be significantly different, about 10% of the total annotated lipid features (Fig. S6B). Therefore, we pooled our female controls to increase statistical power, while proceeding with the Cre and tamoxifen control in males. VIP analysis of lipids most responsible for the separation between genotypes identified phosphatidylcholine and sphingomyelin species to have the highest scores (Fig. 4B). We next performed a relative lipid composition analysis in control and knockout mice of both sexes. We observed that though phosphatidylethanolamine (PE) was the most abundant lipid class across genotypes in both sexes, its relative abundance appeared lower in female *Pdk1*cKO hearts, accompanied with an increase in relative abundance of phosphatidylcholines (Fig. 4C-D, S6C-D). We identified 164 lipid species that were differentially abundant in female hearts and 49 in male hearts, with 33 lipids common to both sexes (Fig. 4E, S6E-F). Majority of the lipids that were upregulated in both male and female belonged to the phosphatidylcholine and sphingomyelin classes (Fig. 4F, S6F). Surprisingly, female *Pdk1*cKO hearts showed a decrease in triacylglycerol species and bulk triacylglycerol abundance (Fig. 4E-F). In contrast, male *Pdk1*cKO hearts showed an increase in a few triacylglycerol species, but bulk triacylglycerol remained unchanged (Fig. S6E,G). Functional lipid analysis^37^ in female *Pdk1*cKO hearts showed a decrease in phospholipid/sphingomyelin, phosphatidylethanolamine/phosphatidylcholine and unsaturated/saturated phosphatidylcholine indices, all indicative of a changing membrane composition and increased membrane rigidity (Fig. 4G). We observed an increase in structural to energetic index, suggestive of increased partitioning towards membrane synthesis (Fig. 4H). No functional indices were significantly modified in male mice. These findings prompted a further examination of the acyl chain composition of phosphatidylcholine and triacylglycerols, classes with the most prominent differences. We observed that 18:1 was the dominant acyl chain in triacylglycerols, while 16:0 dominated in phosphatidylcholines across genotypes and sexes (Fig. S6H-K). In female *Pdk1*cKO hearts, decreases in triacylglycerol-associated acyl chains were accompanied with proportional increases in the same acyl chains within phosphatidylcholines, consistent with a partitioning of acyl chains towards phospholipid synthesis and away from storage (Fig. S6H-I). In male *Pdk1*cKO hearts, this relationship was not observed (Fig. S6J-K). These findings uncover evidence of membrane remodeling, marked by elevated phospholipid acyl chain partitioning and elevated saturated phosphatidylcholines.

**Figure 4.**
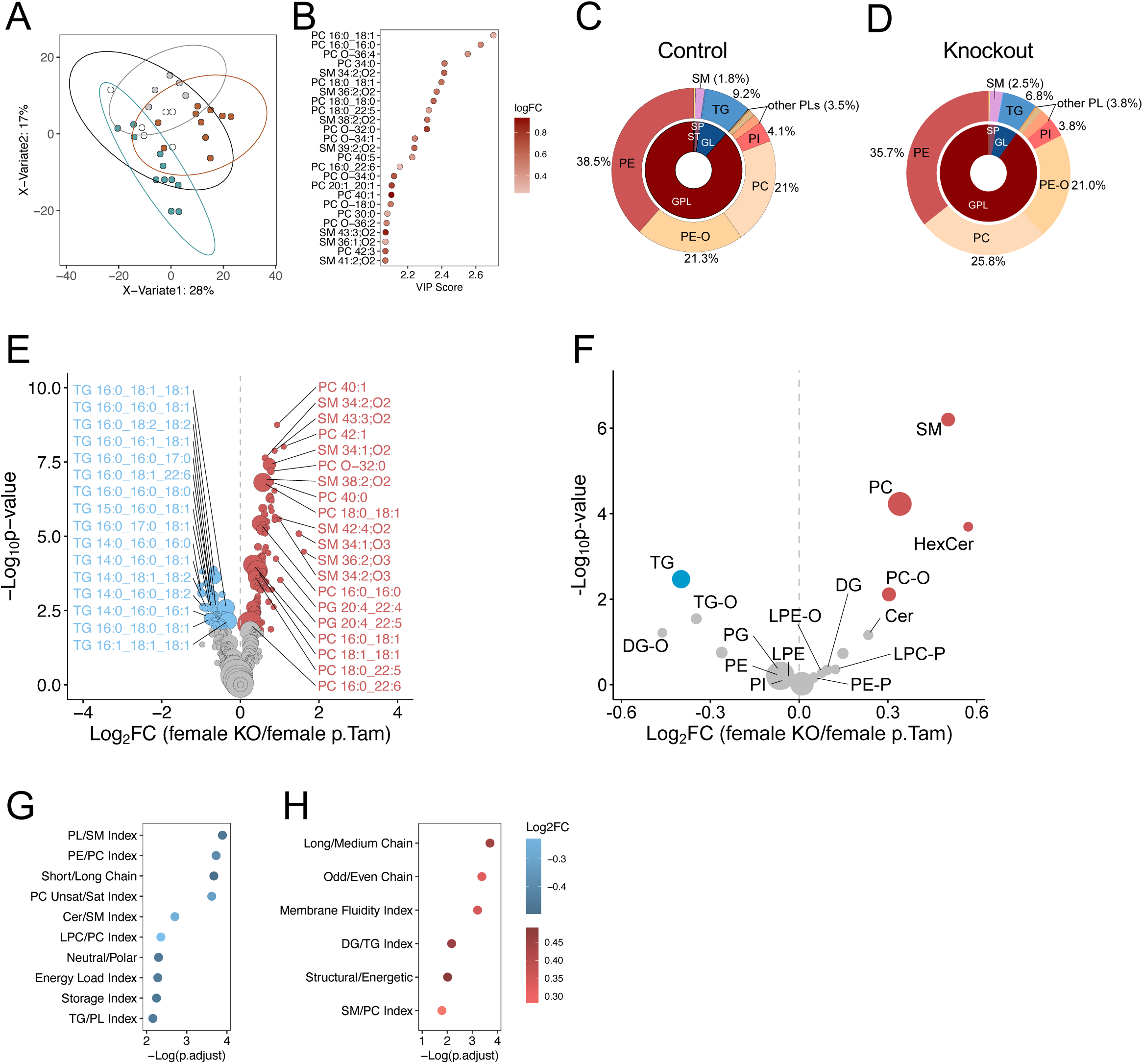
PDK1 reduction alters the cardiac lipidome in female mice hearts. **(A)** Partial least square discrimination analysis of lipid features in mice hearts, 2-weeks post-tamoxifen injection. **(B)** Top 25 lipid with the highest variable importance projection component 1 scores. Red dots are upregulated in female *Pdk1cKO* hearts. Relative lipid composition in female **(C)** control and **(D)** *Pdk1*cKO hearts. Volcano plot depicting differentially abundant **(E)** lipid features and **(F)** lipid classes in female *Pdk1*cKO relative to control. Functional enrichment of female lipidomics dataset using LipidOne platform (v.2.4) depicting **(G)** downregulated and **(H)** upregulated functional indices in female *Pdk1*cKO. Scatter plot visualising phosphosites with a sex-specific response. Gene set enrichment analysis on collapsed phosphoproteins based on the reactome database annotation depicting **(E)** downregulated and **(F)** upregulated pathways in female *Pdk1*cKO hearts. Data were analyzed by Limma. Female pooled Tam control n = 11, male CreTam control n = 5, female *Pdk1*cKO n = 11, male *Pdk1*cKO n = 10.

We next determined how membrane lipid remodeling was related to fatty acid metabolism by quantifying the abundance of acyl carnitines, acyl CoA, carnitine, CoA and free fatty acids using UHPLC-MS/MS metabolomics. We identified 82 metabolites corresponding in these groups. Although most of the annotated features were not significantly different, we observed a downward trend in acyl carnitines, acyl CoA, and free fatty acid features in female *Pdk1*cKO hearts (Fig. 5A). In male mice however, acyl carnitines tended to increase, acyl CoA tended to decrease although to a lesser extent than in males, and fatty acid features showed a general decreased trend (Fig. S7A). We examined the bulk abundances of these metabolite classes and observed a decrease in the free fatty acids and acyl CoA pool in female *Pdk1*cKO hearts (Fig. 5B,C), suggesting a reduced fatty acid availability. Male *Pdk1*cKO hearts also showed a decrease in free fatty acids but no changes to bulk acyl CoA (Fig. S7B-C). Meanwhile, bulk acyl carnitine did not differ in male or female hearts (Fig. 5D, S7D). We examined the ratio of carnitine to acyl carnitine to determine impacts on beta-oxidation and found no differences between control and knockout in both sexes (Fig. 5E, S7E), suggesting no effects on mitochondria acyl transport. Together, these data suggest a reduced fatty acyl availability in female *Pdk1*cKO hearts that may not be attributable to impaired mitochondrial fatty acid import.

**Figure 5.**
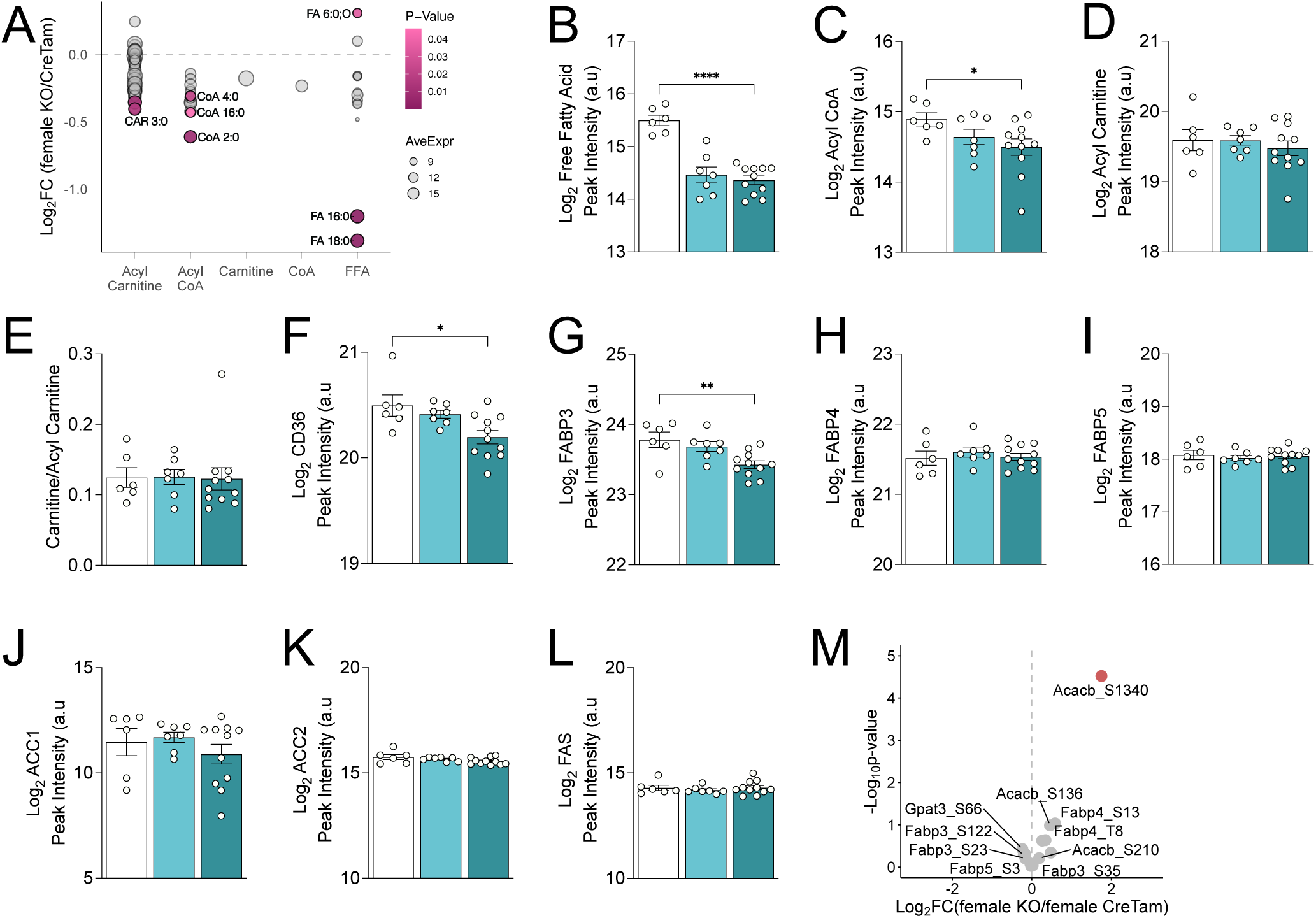
PDK1 reduction decreases fatty acid pools in female mice hearts. **(A)** Fold change of acyl metabolites in *Pdk1*cKO relative to control in female mice hearts, 2-weeks post-tamoxifen injection. UHPLC-MS/MS based abundance of bulk **(B)** free fatty acids **(C)** acyl CoA **(D)** acyl carnitine **(E)** carnitine to acylcarnitine ratio in female mice. Protein abundances of **(F)** CD36; Cluster of Differentiation 36 **(G)** FABP3; Fatty Acid Binding Protein 3 **(H)** FABP4; Fatty Acid Binding Protein 4 **(I)** FABP5; Fatty Acid Binding Protein 5 **(J)** ACC1; Acetyl-CoA Carboxylase 1 **(K)** ACC2; Acetyl-CoA Carboxylase 2 **(L)** FAS; Fatty Acid Synthase. **(M)** Volcano plot depicting differentially phosphorylated lipogenesis phosphosites in female mice. Data are shown as mean ± SEM and were analyzed by either a Limma test (A) or an unpaired two-tailed Welch’s t-test (B-E). Female CreTam control n = 6, female *Pdk1*cHET n = 7, female *Pdk1*cKO n = 11. ^∗^ *p* < 0.05; ^∗∗^ *p* < 0.01; ^∗∗∗^ *p* < 0.001; ∗∗∗∗ *p* < 0.0001.

We tested whether decreased lipogenesis may be responsible for the lowered fatty acyl availability in female *Pdk1*cKO hearts. We used our proteomics dataset to investigate the abundances of lipogenesis proteins. We observed a reduction in fatty acid uptake proteins such as CD36 and the heart-specific FABP3 isoform in female *Pdk1*cKO hearts, suggesting decreased fatty acid uptake (Fig. 5F,G). Other FABP isoforms, the fatty acid synthesis rate-limiting enzyme ACC1/2 and fatty acid synthase FAS were unchanged (Fig. 5H-L). Our phosphoproteomics data showed a significant upregulation in phosphorylation of Acacb_S1340 (ACC2) (Fig. 5M). Though not yet functionally characterized, phosphorylation of ACC1/2 is known to decrease its activity^38^. Male hearts showed no significant changes (Fig. S7F-L). Taken together, our findings indicate decreased fatty acid uptake in female *Pdk1*cKO hearts.

## Discussion

In this study, we deleted *Pdk1* in adult cardiomyocytes using an inducible, cardiomyocyte-specific Cre in high fat diet fed mice of both sexes. Female *Pdk1*cKO hearts had a decrease in left ventricular diameter and volume, but no significant changes in diastolic or systolic function. Multi-omics analysis revealed decreased fatty acid availability and metabolic remodeling encompassing membrane phospholipid composition, energy and lipid metabolism that preceded these anatomical changes.

Mechanistically, multiple lines of proteomic and phosphoproteomic evidence implicate Rho GTPase signaling in the female-specific cardiac phenotype after *Pdk1* knockout. Rho GTPases are molecular switches that promote cytoskeleton re-organization^39^. Cardioprotective roles for RhoA have been proposed based in cardiomyocyte-specific gain- and loss-of-function studies, through mechanisms that could include mitochondrial quality control^40^. In cardiomyocytes, active RhoA binds ROCK to promote cytoskeleton re-organization and contraction^41^. MYPT2 is a downstream phosphorylation target of ROCK that regulates the catalytic activity of myosin light chain phosphatase (MLCP), which subsequently dephosphorylates the myosin regulatory light chain (MLC)^42^. MYPT2 phosphorylation inhibits its activity and results in the inhibition of MLCP. Indeed, Knockout of MYPT2 increases MLC phosphorylation and promotes the existence of myosin in the disordered relaxed state^30^, where it is capable of quickly binding to actin for contraction^43^. Interestingly, MYPT2 knockout hearts showed reduced left ventricular internal diameter at diastole and systole, with elevated fractional shortening^44^, similarly to our finding in female *Pdk1*cKO hearts. Studies have also found that overexpression of MYPT2 desensitizes Ca^2+^ contraction and impairs cardiac function^45^, establishing a potential link between Rho signaling, MYPT2 and Ca^2+^ handling that is evident in female *Pdk1*cKO hearts. It remains to be determined exactly how PDK1 and Rho are linked in a female-specific manner and whether this is related to PDH activity. One potential mechanistic link between PDK1 and Rho signaling is the AMPKα1-mediated inhibitory phosphorylation of RhoA, which occurs in an estradiol-dependent manner^46^. Interestingly, in female *Pdk1*cKO hearts, AMPKα1 is hyperphosphorylated at ser508 (Prkaa1_S508) (Fig. 3C), which has been functionally characterized to inhibit its activity^47^. Inhibition of AMPKα1 could then result in activation of Rho signaling in these female mice. Though, our phosphoproteomics dataset did not annotate other putative targets of AMPKα1 for orthogonal validation, our observation of reduced CD36, the fatty acid uptake protein may point to a diminished effect of AMPK^48^ in female *Pdk1*cKO hearts. How PDK1 deletion causes AMPKα1 hyperphosphorylation at ser508 is unclear, but links between the PDK family and AMPK have been reported. In the liver, PDK4 deficiency decreases ATP levels, which elevates AMPK activity^49^. In our PDK1-deficient hearts, the downregulation of aerobic respiration pathways points to a lower energy profile where we would expect increased, AMPK activity.

Another surprising finding was the differential response between male and female *Pdk1*cKO hearts in PDH activity. Previously, we characterized the impact of PDK1 reduction on PDH activity in a serum starved cardiomyocyte cell line model and observed no changes to PDH activity^16^, in line with previous studies that found no changes to overall PDK activity following PDK1 deletion in male mice hearts^20^. Male *Pdk1*cKO hearts in our study are consistent with previous characterizations, while female *Pdk1*cKO hearts show elevated PDH activity. It is unclear, however, if the overall sex differences observed in our data is a function of the differential response in PDH activity. Our previous studies demonstrated that chronic deletion of PDK1 decreases mitochondrial fatty acid availability and elevates cardiac triacyglycerols in a cardiomyocyte cell line model and high fat diet fed male mice^16^, suggesting a moonlighting function for PDK1 in regulating fatty acid availability. It is plausible that PDK1 regulates fatty acid availability through a mechanism that requires PDH activity in female hearts. PDH determines glucose-derived carbon entry into the citric acid cycle. Therefore, its activity directly dictates the rate of cellular glucose oxidation^50–52^. So, increased PDH activity in female *Pdk1*cKO mice suggests elevated glucose oxidation. The antagonistic nature of the relationship between glucose and fatty acid utilization^53–57^ predicts that elevated glucose oxidation in female *Pdk1*cKO mice hearts may result in decreased fatty acid oxidation dependency. Indeed, the observed decreases in fatty acid uptake proteins and in the abundance of fatty acids are suggestive of a decreased fatty acid oxidation dependency that consequently result in a decreased accumulation of cardiac triacylglycerols. In contrast, male mice hearts with an acute deletion of PDK1, may be more resilient than females by having an unchanged PDH activity and glucose oxidation and consequently no effects on fatty acid metabolism in the interim. Prolonged deletion of PDK1 then may activate the moonlighting functions of PDK1 to decrease fatty acid availability. It is ambiguous what the mechanistic impact of a chronic deletion of PDK1 may have on female mice hearts and present exciting opportunities for future studies.

Aging is characterized by increased left ventricular volume in mice of both sexes^58,59^ and cardiac dysfunction is correlated with age, providing clues that a reduction in internal diameter may be protective. Alternatively, smaller left ventricular diameter has been significantly associated with adverse outcomes in humans^60,61^. Additionally, increased phospholipid saturation, observed in female *Pdk1*cKO hearts, has been associated with the development of cardiac dysfunction^62^, consistent with the idea that PDK1 deficiency may be deleterious on balance. The collective roles of PDK1 in cardiomyopathies justify additional study. Phosphoproteomic analysis of diabetic and ischemic cardiomyopathic patients found dephosphorylation at the PDK1-specific ser232 site on PDH, suggesting reduced PDK1 activity^15^.

In contrast, a recent single-cell transcriptomics study found PDK1 upregulation, though non-significant, in ventricular cardiomyocytes in ischemic cardiomyopathic patients^63^. Additional, cell-type specific proteomic and phospho-proteomic analysis of normal and failing human hearts would help clarify these knowledge gaps.

### Limitations and overall conclusions

Our conclusions, in some cases, are limited by small and uneven sample sizes. In this work, for example, male mice appear to show greater biological variability than female mice. It remains a possibility that our lack of detection of a metabolic or physiological response stems from this variability and may be resolved with larger sample sizes and more prolonged experiments. Additionally, in our study, PDK1 reduction was accompanied with a modest, but significant reduction in PDK2 protein abundance in females although *Pdk2* gene expression was not changed. PDK1 heterodimerizes with PDK2^64^ and knockout of PDK2 in skeletal muscle has resulted in a compensated increase in PDK1^65^. It is unclear if this dimerization is present in female cardiac tissue and the functional implications of it. The modest changes to PDK2 activity may provide an alternative explanation for the sexual dimorphic response on PDH activity.

Despite these caveats, our study reveals a sex-specific cardiomyocyte response to PDK1 loss, potentially mediated through a sex-specific effects on PDH activity and Rho GTPase signaling that link the metabolic remodeling to cardiac structural remodeling. Given the roles of PDK1 in heart failure, our results provide new avenues to modulate cardiomyocyte form and function.

## Methods

### Animals

Animals were housed in ventilated cages within the Modified Barrier Facility at the University of British Columbia with 12-h light:dark cycles. PDK1 floxed mice were obtained from Dr. Eli Zelzer and backcrossed to the C57BL/6J background. *Tg(Myh6-cre/Esr1*)1Jmk^+/-^*mice were obtained from Jackson laboratory (Cat. No 005650) and bred with *Pdk1^fl/fl^* mice to generate *Pdk1^fl/wt^*; *Tg(Myh6- cre/Esr1*)1Jmk^-/-^* and *Pdk1^fl/wt^*; *Tg(Myh6-cre/Esr1*)1Jmk^+/-^* mice. Experimental mice were then generated by crossing *Pdk1^fl/wt^*; *Tg(Myh6-cre/Esr1*)1Jmk^-/-^*mice with *Pdk1^fl/wt^*; *Tg(Myh6- cre/Esr1*)1Jmk^+/-^* mice and weaned at 3 weeks on a high fat diet 20% kcal sucrose, and 60% kcal fat (D12492i, Research Diets). At 8-10 weeks, experimental mice were doubly administered an intraperitoneal injection of 20 mg/mL tamoxifen (Cat. No. T5648, Sigma-Aldrich, Oakville, ON, CA) dissolved in corn oil (Cat. No. C8267, Sigma-Aldrich, Oakville, ON, CA) or just corn oil at a dose of 1 mg/20 g body weight, 48 h apart. Two weeks post injection, mice were fasted for 4 h and euthanized by CO_2_ followed by cervical dislocation in accordance with the Animal Care guidelines. At this time, mice were weighed, and tissues were rapidly harvested and flash frozen in liquid nitrogen. All procedures were approved by the University of British Columbia Animal Care and Use Committee and were performed in accordance with Canadian Council on Animal Care Guidelines (Protocol No. A21- 0135).

### Gene Expression Analysis

Frozen tissues were ground in a liquid nitrogen pre-cooled mortar and pestle. Total RNA was extracted from approximately 20 mg of ground tissue using the Qiagen RNeasy extraction kit (Cat. No. 74104, Qiagen, Hilden, DE), after which cDNA was synthesized using the qScript cDNA synthesis kit (Cat. No. 101414-100, Quanta Biosciences, Beverley, US) and High-Capacity cDNA Reverse Transcription Kit (Cat. No. 4368813, ThermoFisher Scientific, Burnaby, BC, CA). Gene expression was measured with using real time qPCR and Taqman gene expression assay (ThermoFisher Scientific, Burnaby, BC, CA). β-actin (*Actr3b*, Cat. No. Mm01134842_m1, ThermoFisher Scientific, Burnaby, BC, CA) was used as a house keeping gene. Cycle threshold (Ct) values of genes (*Pdk1*, Cat. No. Mm00554300_m1; *Pdk2*, Cat. No. Mm00446681_m1; *Pdk3*, Cat. No. Mm00455220_m1; *Pdk4*, Cat. No. Mm01166879_m1, ThermoFisher Scientific, Burnaby, BC, CA) were subtracted from Ct values of β-actin to obtain relative expression to β-actin (ΔCt), which was then converted to linear scale (2^-ΔCt^).

### PDH Activity Assay

Frozen tissues were ground in a liquid nitrogen pre-cooled mortar and pestle. Ground tissues were homogenized in 100 μL of ice-cold assay buffer from the pyruvate dehydrogenase activity kit (Cat. No. MAK183, Sigma-Aldrich, Oakville, ON, CA), with 40-60 passages in a Dounce homogenizer. Samples were incubated on ice for 10 min and then centrifuged at 10, 000 xg for 5 min at 4 °C. 2.7 M ammonium sulphate was added to the supernatant, incubated on ice for 20 min and centrifuged at max speed for 5 min at 4 °C. The resulting supernatant was discarded, and pellet resuspended in 200 μL of assay buffer. Enzyme activity was then measured per the manufacturer’s instructions.

### Echocardiography

Cardiac function was determined using a Vevo 2100 imaging system (Fujifilm VisualSonics, Inc., Toronto, Canada). Before imaging, mice were anesthetized in inhaled isoflurane (2–3% for induction and 1–2% for maintenance) during echocardiographic assessment and hair removed from the ventral thorax region using a depilatory cream. M-mode images were acquired from parasternal long-axis and short-axis views at the mid-papillary muscle level to assess cardiac structure and function. Images were digitally recorded and analyzed offline using Vevo LAB software (Fujifilm VisualSonics, Inc., Toronto, Canada) by an investigator blinded to the experimental groups. Left ventricular (LV) internal diameter, LV posterior wall thickness, and interventricular septal (IVS) thickness were measured during systole and diastole. Functional parameters including LV mass, ejection fraction, fractional shortening, heart rate, and indices of diastolic function were also calculated from M-mode recordings. Mitral inflow and tissue Doppler measurements were also obtained to evaluate diastolic function.

### Proteomics and Phosphoproteomics

Cryopulverized mouse cardiac tissue was extracted in tissue lysis buffer (5% SDS, 100 mM TRIS pH 8.5, 10 mM TCEP) at 95°C for 10 minutes with rotational agitation at 1800 rpm using an Eppendorf ThermoMixer C. Lysates were clarified by centrifugation at 21,000 x g for 1 minute, and the supernatant was transferred to new microcentrifuge tubes. Protein concentrations were determined using a reducing agent compatible bicinchoninic acid assay kit (Thermo Fisher Scientific / Pierce). The remaining sample was alkylated in the presence of 40 mM iodoacetamide for 20 minutes in the dark. An equivalent of 50 µg of protein was proteolytically digested with sequencing grade trypsin (Promega) at 37°C overnight (16 hours) using S-TRAP micro cartridges according to the vendor protocol (Protifi LLC). 1% of the sample was reserve for global proteome profiling by DIA-PASEF MS (described below) while the remaining digested peptides were lyophilized to dryness and solubilized in 80% acetonitrile with 1% TFA. Phosphopeptides were then enriched using zirconium immobilized metal affinity chromatography magnetic nanoparticles (Zr-IMAC HP, Resyn Biosciences). After enrichment, phosphopeptides were eluted in 1% ammonium hydroxide, then neutralized by the addition of 5% TFA and vacuum concentrated to dryness prior to rehydration in 0.1% formic acid prior to loading onto evotips. For global proteomics, an equivalent of 200 nanograms per sample was loaded onto Evotip Pure tips (Evosep) according to the manufacturer protocol and analyzed using an Evosep One LC and a Bruker timsTOF HT mass spectrometer operated in DIA-PASEF mode using an empirically optimized isolation scheme using the py_diAID tool ^66^ using 12 PASEF ramps split with 24 MS/MS windows, and had an estimated cycle time of 1.2 s. Precusor isolation was between an 100 – 1700 m/z and an ion mobility range of 0.7 – 1.4 1/k0 with ramp and accumulations times set to 100 ms. Collision energy was scaled based on precursor ion mobility from 20 – 59 eV between 0.6 and 1.6 1/k0 respectively. DIA-PASEF data was analyzed using DIA-NN (2.3.2) using an in silico predicted spectral library generated by DIA-NN, based on the canonical human reference proteome (UP000005640, containing 20,464 protein sequences) and the default application settings (MBR enabled, protein inference selected, proteotypicity based on gene sequences, cross run normalization based on retention time). Protein group level quantitation (report.pg_matrix.tsv) was used for downstream statistical analysis.

For phosphoproteomics analysis, DDA-PASEF was conducted using the same instrumentation as noted above with MS survey scan (100 – 1800 m/z) and 10 PASEF ramps isolating the most abundant precursors for MS/MS fragmentation, with active exclusion of previously triggered precursors set to 0.4 minutes. PASEF intensity threshold and targets were set to 2500 and 20000 respectively. The ion mobility window was set between 0.7 and 1.4 1/K0. The CaptiveSpray source voltage was set to 1.6 kV.

Collision energy was scaled from 20 – 59 eV between 0.6 and 1.6 1/K0. Raw data was searched, and label free quantitation was performed using Fragpipe version ^67^. Database searching used reference human proteome sequences from Uniprot (Uniprot, UP000005640). Phosphorylation of serine (S), threonine (T), and tyrosine residues was defined as a variable modification. Features without quantitation in at least 70% of a given experimental condition were removed from the analysis. Differential expression analysis and visualization was conducted using limma in R. Significance was based on a Benjamini-Hochberg FDR adjusted *p* < 0.05. Missing values were imputed using the perseus-type scheme in Fragpipe analyst. Pathway enrichment was performed using a gene-set enrichment analysis, where proteins were ranked by the *p* value and the sign of Log_2_ fold change to determine directionality, and pathways annotated using the reactome database.

### Cytoscape network representation

Since the female proteomics dataset had >100 differentially abundant proteins, a rank score was obtained by multiplying the absolute value of the log_2_(fold change) by the negative logarithm of the adjusted *p* value (-log_10_(adjusted *p* value)). The dataset was then filtered for the top 100 hits based on the highest rank score. No additional filtration threshold was applied to the male proteomics dataset since it had <100 differentially abundant proteins. Proteomics datasets were then merged with the sex-appropriate phosphoproteomics dataset and queried for a protein-protein interaction network with a medium confidence score (STRING functional score 0.4). Subcellular location was assigned as the location with the highest score (0-5) obtained in STRING. When there was a tie, Uniprot was queried to determine the most appropriate location.

### Lipidomics

Untargeted lipidomics was performed on a Bruker timsTOF Pro2 high-resolution mass spectrometer (Bruker Daltonics, Bremen, Germany) coupled to an Elute UHPLC system (Bruker Daltonics). Dried lipid extracts were reconstituted in 250 µL of acetonitrile:isopropanol (70:30, v/v) containing 0.1 ppm of 12- [[(cyclohexylamino)carbonyl]amino]-dodecanoic acid (CUDA; Cayman Chemicals) and Equisplash™ (Cat No. 330731, Avanti Research, Alabama, US) as internal standards, vortexed for 10 min and centrifuged at 14,000 rpm for 10 min prior to injection. Aliquots of 20 µL from every sample were pooled to generate pooled quality-control (QC) samples, which were injected every 12 samples together with extraction blanks to monitor instrument performance. Sample order was randomized to minimize batch-related artifacts.

Lipid separation was achieved on an Acquity UPLC CSH C18 column (130 Å, 1.7 µm, 2.1 × 100 mm; Waters, Milford, MA), using a method adapted from Cajka et al.^68^, as previously described ^16^. Briefly, mobile phase A was acetonitrile:water (60:40, v/v) and mobile phase B was isopropanol:acetonitrile (90:10, v/v); both phases were supplemented with 0.1% (v/v) formic acid and 10 mM ammonium formate. A multi-step gradient from 15% to 99% B was delivered over 17 min as follows: 0 min, 15% B; 0–2 min, 30% B; 2–2.5 min, 50% B; 2.5–12 min, 80% B; 12–12.5 min, 99% B; 12.5–13.5 min, 99% B; 13.5–13.7 min, 15% B; 13.7–17 min, 15% B. The flow rate was 0.5 mL min^−1^, the column was maintained at 65 °C, the autosampler at 4 °C and the injection volume was 2 µL.

Mass spectra were acquired over a mass range of 100–2,000 *m/z* in data-dependent acquisition mode with parallel accumulation–serial fragmentation (ddaPASEF). TIMS was operated with a 100 ms accumulation time and a 100 ms ramp time over a mobility (1/K_0_) range of 0.55–1.90 V·s cm^−2^, giving an effective duty cycle of 100% and a resulting spectra rate of 9.41 Hz. Each ddaPASEF cycle included two MS/MS scans and targeted three precursors per TIMS ramp, for a total cycle time of 0.5 s. Active exclusion was enabled with an exclusion-release time of 0.10 min, and the ion charge control (ICC) target was set to 7.5 × 10^6^ counts. Collision energy was ramped linearly as a function of ion mobility: in positive ion mode (ESI^+^), the collision energy was increased from 31.92 eV at 1/K_0_ = 0.80 V·s cm^−2^ to 51.24 eV at 1/K_0_ = 1.60 V·s cm^−2^ (values at 0.65, 0.80, 1.00, 1.20, 1.40 and 1.60 V·s cm^−2^ were 42.00, 31.92, 36.96, 42.00, 47.04 and 51.24 eV, respectively); in negative ion mode (ESI^−^), the same mobility-dependent ramp was applied with collision energies of −42.00, −31.92, −36.96, −42.00, −47.04 and −51.24 eV at the corresponding mobility values.

Source and transfer conditions were optimized independently for each polarity. In ESI^+^, the capillary voltage was set to 4,500 V with an end-plate offset of −500 V, nebulizer pressure 2.2 bar, dry-gas flow 9.0 L min^−1^ and dry-gas temperature 220 °C; the collision cell RF was 1,100 Vpp and the transfer time was 70 µs. In ESI^−^, the capillary voltage was set to −4,500 V with an end-plate offset of −500 V, nebulizer pressure 3.0 bar, dry-gas flow 6.0 L min^−1^ and dry-gas temperature 230 °C; the collision cell RF was 1,000 Vpp and the transfer time was 50 µs. Internal mass calibration was performed in every run by injecting 10 µL of 10 mM sodium formate through a six-port diverter valve at the beginning of each run, yielding an average mass error below 2 ppm.

Lipid features with MS/MS were then annotated using Metaboscape (Bruker Daltonics, Bremen, Germany) and involved peak picking, alignment, and database searching. Features with >25% relative standard deviation across QC samples were filtered out of the data. Lipid classes were normalized by internal standard peak intensity. Duplicate features across positive and negative modes were processed by selecting the feature with the lower coefficient of variation across QC samples. Additionally, features with the highest sample peak intensity <5x blank intensity was regarded as not meeting the quantification threshold. The dataset was then transformed to log_2_ scale to normalize the data distribution before statistical tests were performed. Additionally, it was also analyzed on the LipidOne (v 2.4) platform for biological interpretation ^69^.

### Untargeted polar-metabolite profiling by reversed-phase LC–MS/MS

The polar (aqueous) fraction was analysed by reversed-phase LC–MS/MS using an Inertsil Ph-3 UHPLC column as previously described ^70^, complementing the HILIC analysis described below. To track method performance, the extraction solvent was spiked with the internal standards methionine-d3, ferulic acid-d3, and caffeine-^13^C_3_ at 1 ppm. The dried polar fraction was resuspended in 250 µL of aqueous methanol (50%, v/v), centrifuged at 14,000 × *g* for 10 min, and 200 µL of the clear supernatant was transferred to an LC-MS vial. Pooled QC samples were generated by combining 20 µL from each sample; QCs and extraction blanks were injected every 12 samples, and the sample order was fully randomized to minimize batch-related artifacts.

Samples were analyzed on a Bruker Impact II QTOF high-resolution mass spectrometer (Bruker Daltonics) coupled to a Vanquish Horizon UHPLC system (Thermo Fisher Scientific) as previously described ^70^ with some modifications. Separation was achieved on an Inertsil Ph-3 UHPLC column (2 µm, 150 × 2.1 mm; GL Sciences) fitted with a Ph-3 guard column (2 µm, 2.1 × 10 mm). Mobile phase A was water containing 0.1% (v/v) formic acid, and mobile phase B was methanol containing 0.1% (v/v) formic acid. A multi-step gradient from 5% to 99% B was delivered over 18 min as follows: 0 min, 5% B; 0–1 min, 5% B; 1–8 min, 35% B; 8–10.5 min, 99% B; 10.5–14 min, 99% B; 14–14.5 min, 5% B; 14.5–18 min, 5% B. The column temperature was set to 55 °C, the autosampler was held at 4 °C and the flow rate was 0.3 mL min^−1^. Injection volumes were 2 µL in positive ion mode and 3 µL in negative ion mode. Data-dependent acquisitions were performed in both positive (ESI^+^) and negative (ESI^−^) ionization modes. For ESI^+^, the mass spectrometer settings were: capillary voltage 4,500 V, nebulizer pressure 2.0 bar, dry-gas flow 9 L min^−1^, dry-gas temperature 220 °C, mass scan range 60–1,300 *m/z*, spectral acquisition rate 3 Hz and total cycle time 0.6 s. Collision energy of 20 V was ramped from 100% to 250% across each MS/MS scan to obtain comprehensive fragment-ion information. For ESI^−^, the capillary voltage was set to −3,500 V and all other source parameters were held constant. To ensure high mass accuracy, internal calibration was performed in each analytical run by injecting 10 µL of 10 mM sodium formate via a six-port diverter valve from 0 to 0.15 min, yielding an average mass error below 2 ppm.

### Untargeted polar-metabolite profiling by HILIC LC–MS/MS

The polar fraction was also analysed by Hydrophilic Interaction Liquid Chromatography (HILIC) to retain highly polar species, that are poorly retained by reversed-phase chromatography. Samples were analyzed on a Bruker Impact II QTOF mass spectrometer (Bruker Daltonics) coupled to a Vanquish UHPLC system (Thermo Fisher Scientific).

Chromatographic separation was performed on an InfinityLab Poroshell 120 HILIC-Z column (2.7 µm, 150 × 2.1 mm; Agilent Technologies, Santa Clara, CA). Separation was achieved on an InfinityLab Poroshell 120 HILIC-Z column (2.7 µm, 150 × 2.1 mm). Mobile phases consisted of A water with 0.1% formic acid, 10 mM ammonium acetate and 5 µM medronic acid, and B acetonitrile:water (90:10, v/v) with identical additives. The gradient was: 0–2 min, 90% B; 2–7 min, 48% B; 7–7.2 min, 40% B; 7.2–7.8 min, 40% B; 7.8–8.2 min, 90% B; 8.2–12.5 min, 90% B. Column temperature was 30 °C, autosampler 4 °C, and the flow rate was 0.3 mL min^−1^.

QC samples were injected every 12 samples to monitor system stability. Data-dependent MS/MS spectra were collected in both ESI^−^ and ESI^+^ modes. In ESI^−^, the capillary voltage was −3,800 V, nebulizer pressure 2.0 bar, dry-gas flow 9 L min^−1^, dry-gas temperature 220 °C, mass scan range 100– 1,500 *m/z*, spectral acquisition rate 3 Hz and cycle time 0.7 s; collision energy was ramped from 20 V across 100–250% of each MS/MS scan. Quadrupole ion energy was set to 4 eV with a pre-pulse storage time of 7 µs; collision RF and transfer time were ramped from 250 Vpp/40 µs (15% of the cycle) to 850 Vpp/115 µs (85% of the cycle). In ESI^+^, the capillary voltage was set to 4,500 V with all other parameters as above. Internal calibration was performed in each run by injecting 10 µL of 10 mM sodium formate via a six-port diverter valve at the start of each acquisition.

### Data processing and metabolite annotation

Raw LC–HRMS/MS data from the reversed-phase Inertsil Ph-3 polar-metabolite method and the HILIC method were processed with Progenesis QI v.3.0.7600.27622 (Nonlinear Dynamics/Waters) using the METLIN plugin v.1.0.7642.33805 as previously described ^70,71^. Processing comprised alignment, deconvolution, peak picking, normalization and database searching. Normalization was carried out using the built-in “all-compounds” algorithm of Progenesis QI. Molecular features were filtered using the following criteria before downstream analysis: (1) the presence of acquired MS/MS spectra (features without MS/MS were excluded); (2) a coefficient of variation (CV) < 25% across the pooled QC injections; (3) a signal intensity in study samples at least five-fold higher than in extraction blanks; and (4) a mass error below 5 ppm, with average annotation mass error below 2.0 ppm.

Annotations were assigned in accordance with the Metabolomics Standards Initiative (MSI) reporting criteria ^72^. Level 1 (L1) annotations required retention-time and MS/MS spectral match to authentic standards analysed under identical conditions, including an *in-house* library built from the Mass Spectrometry Metabolite Library of Standards (MSMLS; IROA Technologies) as previously described ^70^. Level 2 (L2) annotations were based on experimental MS/MS matching against METLIN, the Human Metabolome Database (HMDB), MassBank of North America (MoNA; https://mona.fiehnlab.ucdavis.edu/) and GNPS. A Progenesis QI score ≥ 40 was used as the minimum threshold for L2 candidate acceptance. When a compound was detected in both polarities, the ion with the lower CV on QCs was kept for downstream analysis.

### Statistical Analysis

Statistical analyses and data presentation for were carried out using GraphPad Prism 9 (Graphpad Software, San Diego, CA, USA) or R (v 4.1.1). Two-tailed Welch’s *t*-tests were performed for pairwise comparisons. For omics data, linear models for microarray (LIMMA-VOOM) R package ^73^ was used in assessing differential abundance. For all statistical analyses, differences were considered significant if the (adjusted using the Benjamini-Hochberg method) *p* value was less than 0.05: ^∗^ p< 0.05; ^∗∗^ p< 0.01; ^∗∗∗^ p< 0.001; ∗∗∗∗ p<0.0001. Data were presented as means ± SEM with individual data points from biological replicates, unless otherwise indicated in the figure legends.

## Author contributions

M.G.A. designed experiments, performed experiments, analyzed data, and wrote the manuscript. S.P. designed experiments, performed experiments, analyzed data. J.J.H., X.H., A.B. performed experiments. H.H.C. analyzed data. S-Y.C. performed experiments, analyzed data. V.R.R., T.G., A.A.M. designed experiments, performed experiments, analyzed data. B.R., L.J.F., E.J.R., C.H.B. analyzed data, provided critical resources. J.D.J. designed experiments, analyzed data, edited the manuscript, provided resources, and is the guarantor of this work.

## Acknowledgements

We thank members of the Johnson lab for their supporting efforts in the completion of this study. We acknowledge Dr. Eli Zelzer for his generosity in sharing the *Pdk1*-floxed mice, Bahira Hussein for her administrative help with the echocardiological experiments, Dr. Alison McAfee for her ingenuity of creating a missingness test, and the excellent animal staff and technicians at the Modified Barrier Facility at UBC.

