## Supplemental figures for "Cardiomyocyte pyruvate dehydrogenase kinase 1 knockout activates Rho-mediated remodeling and decreases fatty acyl availability in female mice"

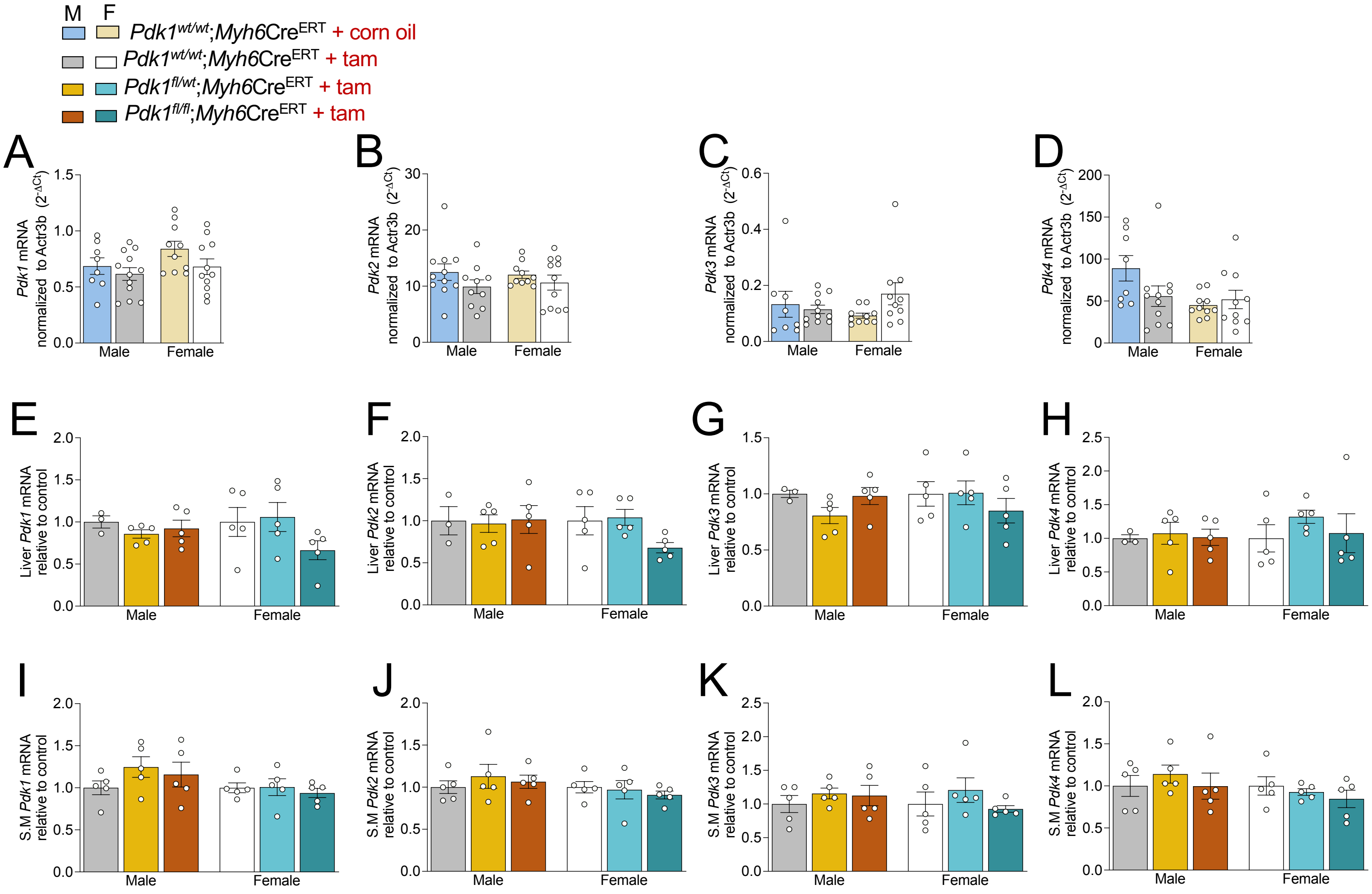


**Fig S1: Model Validation. (A)** *Pdk1*, **(B)** *Pdk2*, **(C)** *Pdk3*, **(D)** and *Pdk4* gene expression in mice hearts, 2 weeks post tamoxifen injection. *Pdk* isoform gene expression in **(E-H)** liver and **(I-L)** skeletal muscle. Data are shown as mean ± SEM and were analyzed by 2-way ANOVA followed by a post-hoc Tukey multiple comparison test. $^{*}\ p < 0.05$; $^{**}\ p < 0.01$; $^{***}\ p < 0.001$; $^{****}\ p < 0.0001$

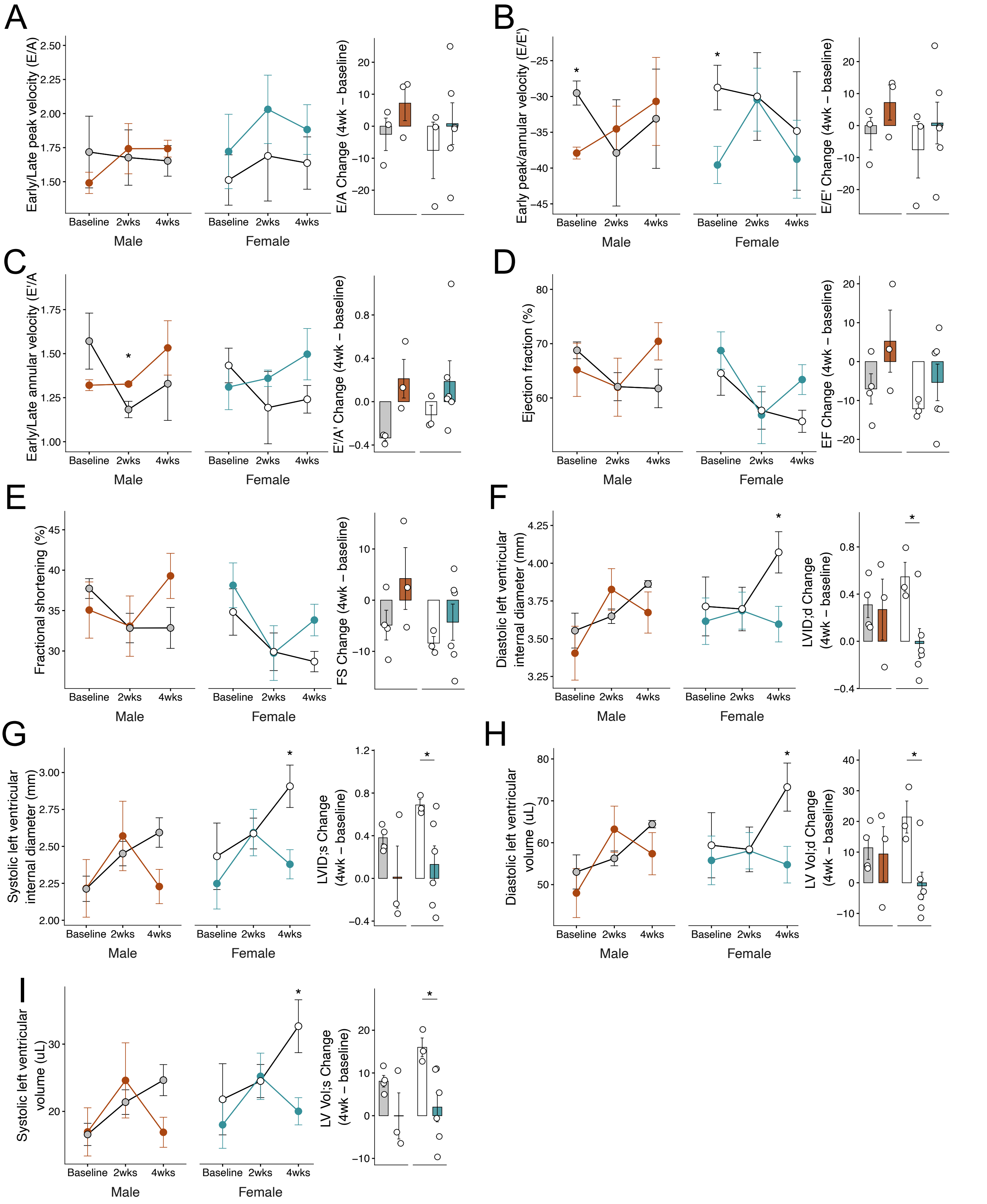


**Fig S2: Effects of PDK1 reduction on cardiac function.** **(A)** Early to late peak velocity (E/A), **(B)** early peak velocity to early annular velocity (E/E'), **(C)** early annular velocity to late annular velocity (E'/A'), **(D)** ejection fraction, **(E)** fractional shortening, **(F)** diastolic left internal diameter, **(G)** systolic left internal left diameter, **(H)** diastolic left ventricular volume, and **(I)** systolic left ventricular volume were assessed in mice before tamoxifen injection (baseline), 2 weeks and 4 weeks post-tamoxifen injection. Data are shown as mean ± SEM and were analyzed by an unpaired two-tailed Student's t-test. * $p < 0.05$; ** $p < 0.01$; *** $p < 0.001$; **** $p < 0.0001$.

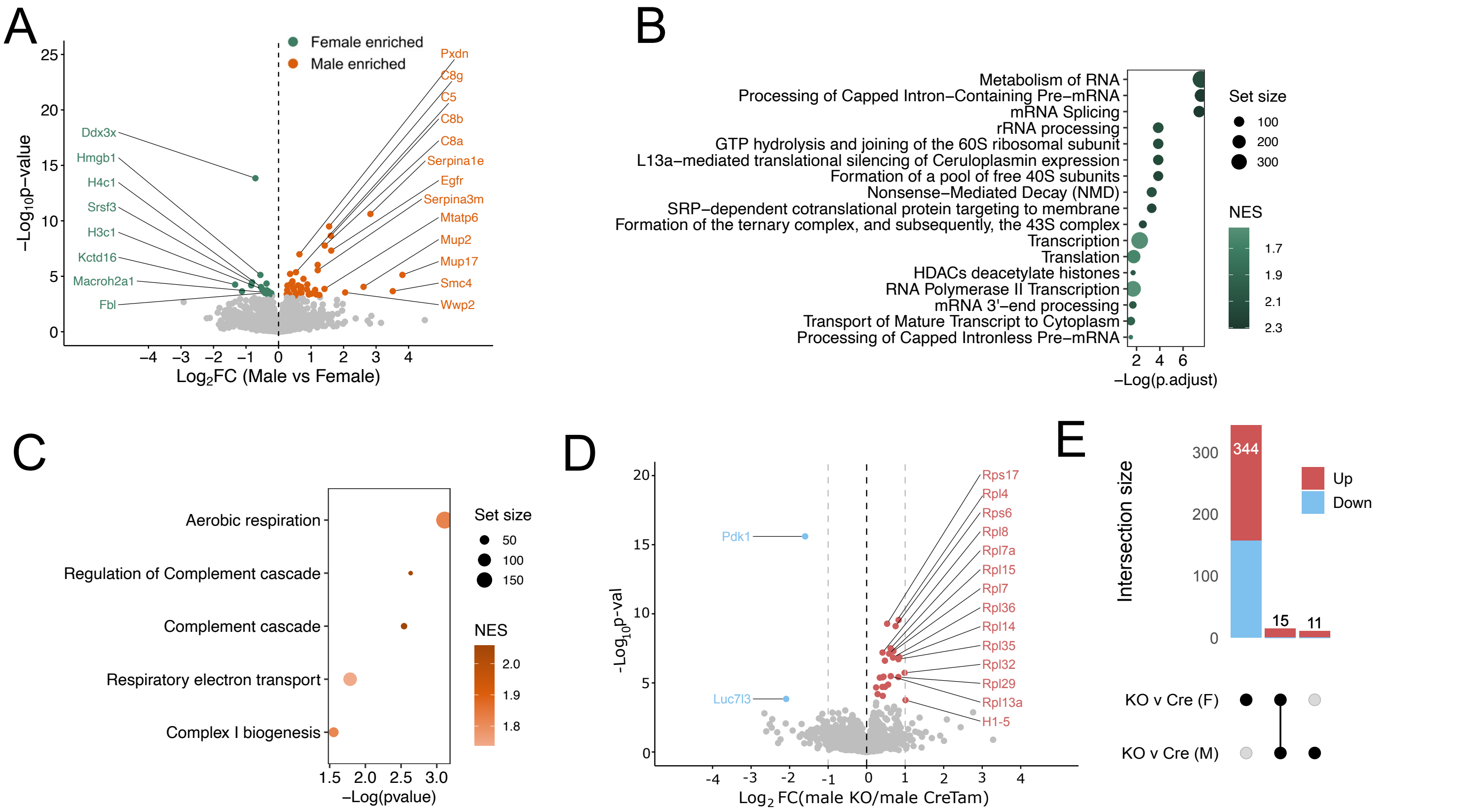


**Fig S3: Effect of biological sex and PDK1 reduction on the cardiac proteome. (A)** Volcano plot depicting differentially enriched proteins in male and female controls. Gene set enrichment analysis based on the reactome database annotation depicting enriched pathways in **(B)** female and **(C)** male control hearts. **(D)** Volcano plot depicting differentially abundant proteins in male knockouts relative to control. **(E)** UpSet plot visualizing the intersections in differential analysis from male and female datasets. Data were analyzed using Limma with a Benjamini-Hochberg method for multiple testing correction and an adjusted $p < 0.05$. Female CreTam control n = 6, male CreTam control n = 5, female *Pdk1*cKO n = 11, male *Pdk1*cKO n = 10.

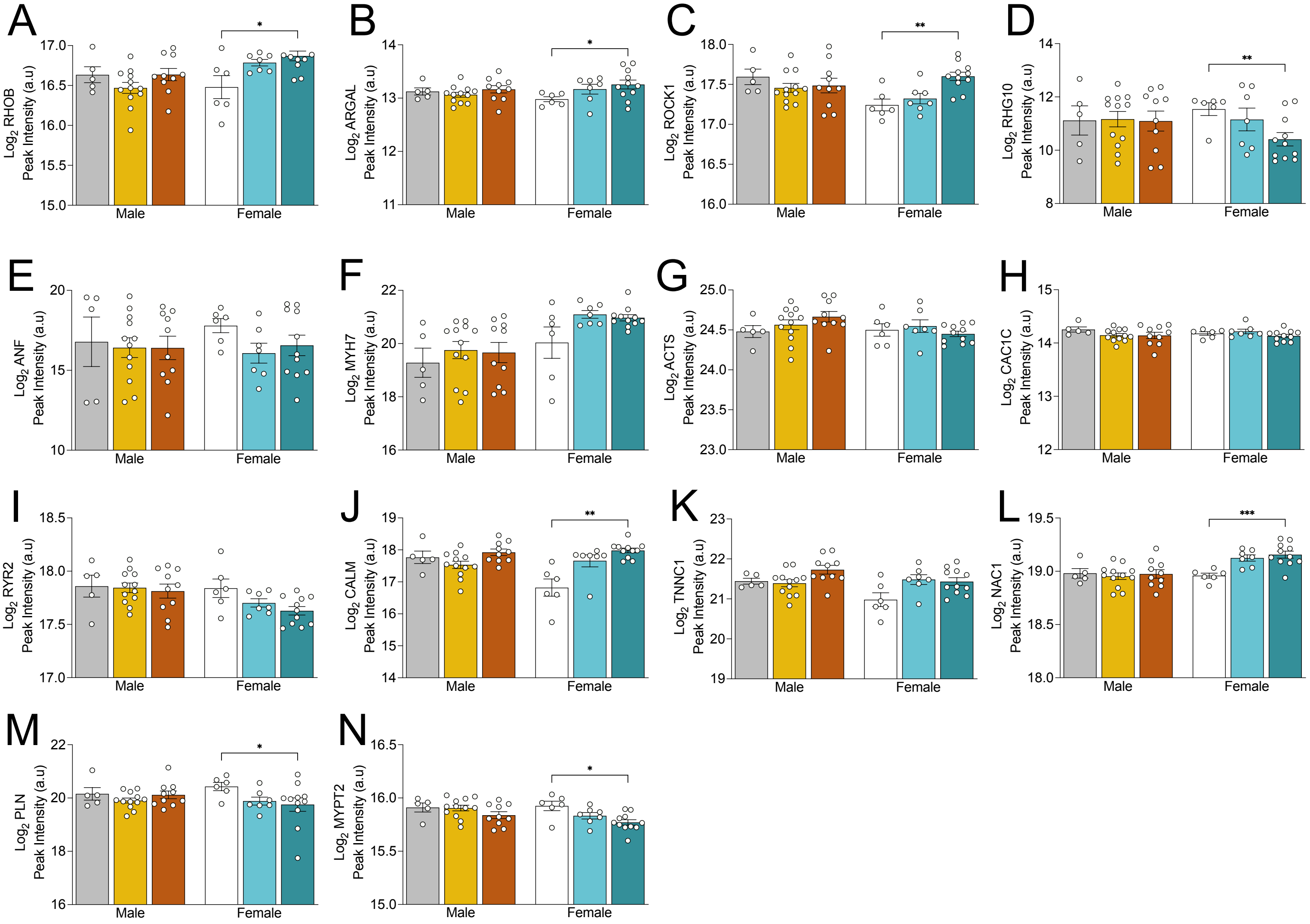


**Fig S4: Evaluation of Rho signaling, cardiac hypertrophic markers and calcium handling proteins.** Protein abundances of **(A)** RHOB; Rho-related GTP-binding protein RhoB **(B)** ARGAL; Rho Guanine Nucleotide Exchange Factor 10-like protein **(C)** ROCK1; Rho-associated Protein Kinase 1 **(D)** RHG10; Rho GTPase-activating protein 10 **(E)** ANF; Atrial Natriuretic Factor **(F)** MYH7; Myosin Heavy Chain 7 **(G)** ACTS; Actin, Alpha Skeletal Muscle **(H)** CAC1C; Voltage-dependent L-type Calcium Channel **(I)** RYR2; Ryanodine Receptor 2 **(J)** CALM; Calmodulin-1 **(K)** TNNC1; Troponin C **(L)** NAC1; Sodium/Calcium Exchanger 1 **(M)** PLN; Phospholamban **(N)** MYPT2; Protein phosphatase 1 regulatory subunit 12B . Data are shown as mean ± SEM and were analyzed using an unpaired two-tailed Welch's t-test (B-E). * $p < 0.05$; ** $p < 0.01$; *** $p < 0.001$; **** $p < 0.0001$.

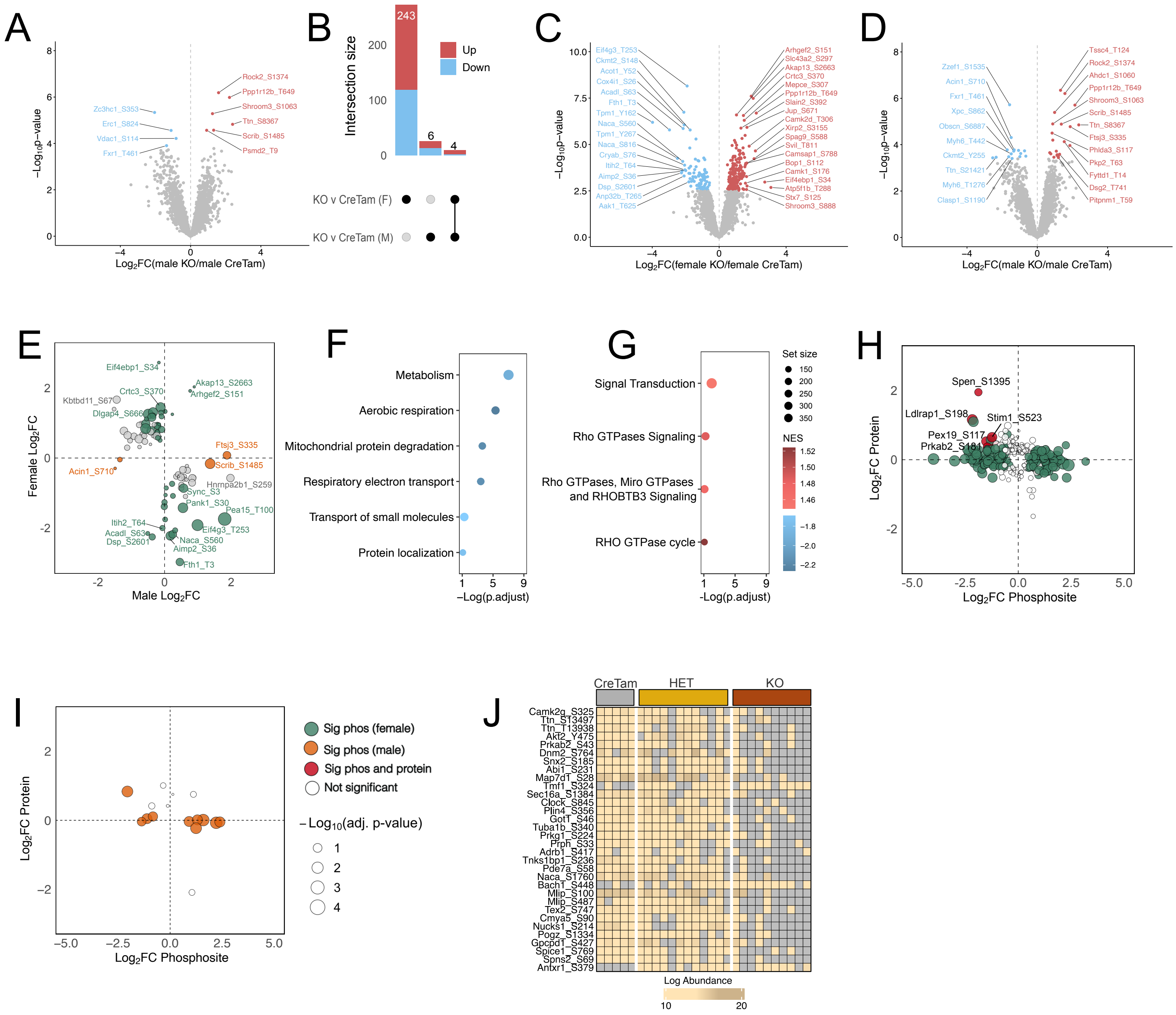


**Fig S5: Effect of PDK1 reduction on the cardiac phosphoproteome. (A)** Volcano plot depicting differentially abundant protein-normalized phosphosites in male *Pdk1*cKO relative to control, 2-weeks post-tamoxifen injection. **(B)** UpSet plot visualizing the intersections in differential analysis from male and female protein-normalized phosphorylation datasets. Volcano plot depicting differentially abundant non-protein normalized phosphosites in **(C)** females and **(D)** males. **(E)** Scatter plot visualising non-protein normalized phosphosites with a sex-specific response to PDK1 reduction. Gene set enrichment analysis on collapsed non-protein normalized phosphoproteins based on the reactome database annotation depicting pathways with **(F)** decreased **(G)** and increased phosphorylation signals in female *Pdk1*cKO hearts. Scatter plot visualizing the fold change across differentially phosphorylated phosphosites and corresponding protein abundance in **(H)** female and **(I)** males. **(J)** Heatmap of phosphosites with significant missingness in male mice evaluated by Barnard's unconditional exact test (nominal p.value < 0.05). Data were analyzed by Limma. Female CreTam control n = 6, male CreTam control n = 5, female *Pdk1*cKO n = 11, male *Pdk1*cKO n = 10.

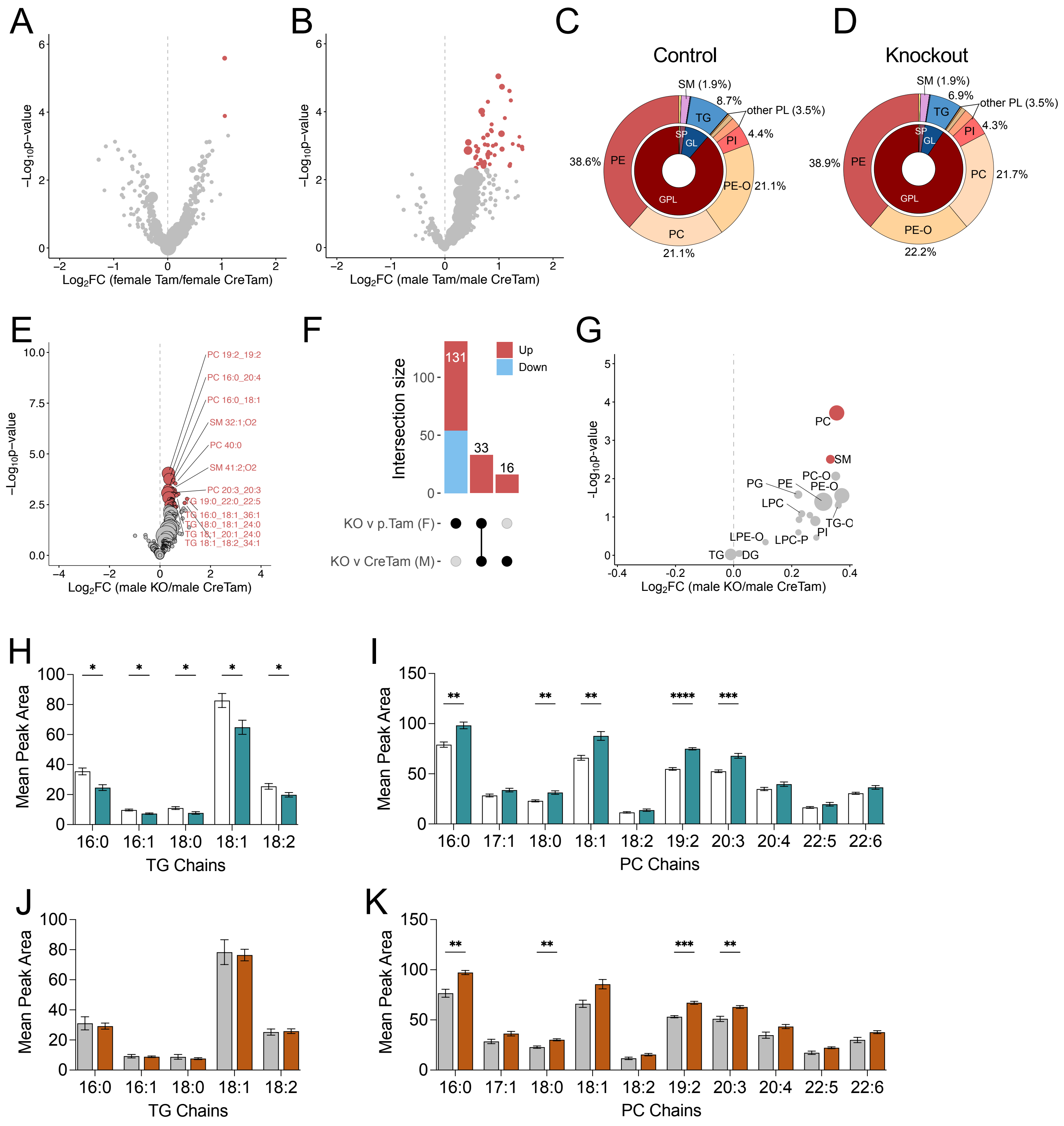


**Fig S6: Effect of PDK1 reduction on the cardiac lipidome.** Volcano plot depicting differentially abundant lipids between tamoxifen (no Cre) control and Cre + tamoxifen control in **(A)** female and **(B)** male mice. Relative lipid composition in male **(C)** CreTam control and **(D)** *Pdk1*cKO hearts. **(E)** Volcano plot depicting differentially abundant lipid features in male knockouts relative to CreTam control. **(F)** UpSet plot visualizing the intersections in differential analysis from male and female datasets. **(G)** Volcano plot depicting differentially abundant lipid features in male *Pdk1*cKO relative to CreTam control. Acyl chain analysis in **(H, J)** triacylglycerols and **(I, K)** phosphatidylcholines in female **(H, I)** and male **(J, K)** using the LipidOne platform (v2.4). Data were analyzed by Limma. Female CreTam control n = 6, male CreTam control n = 5, female *Pdk1*cKO n = 11, male *Pdk1*cKO n = 10.

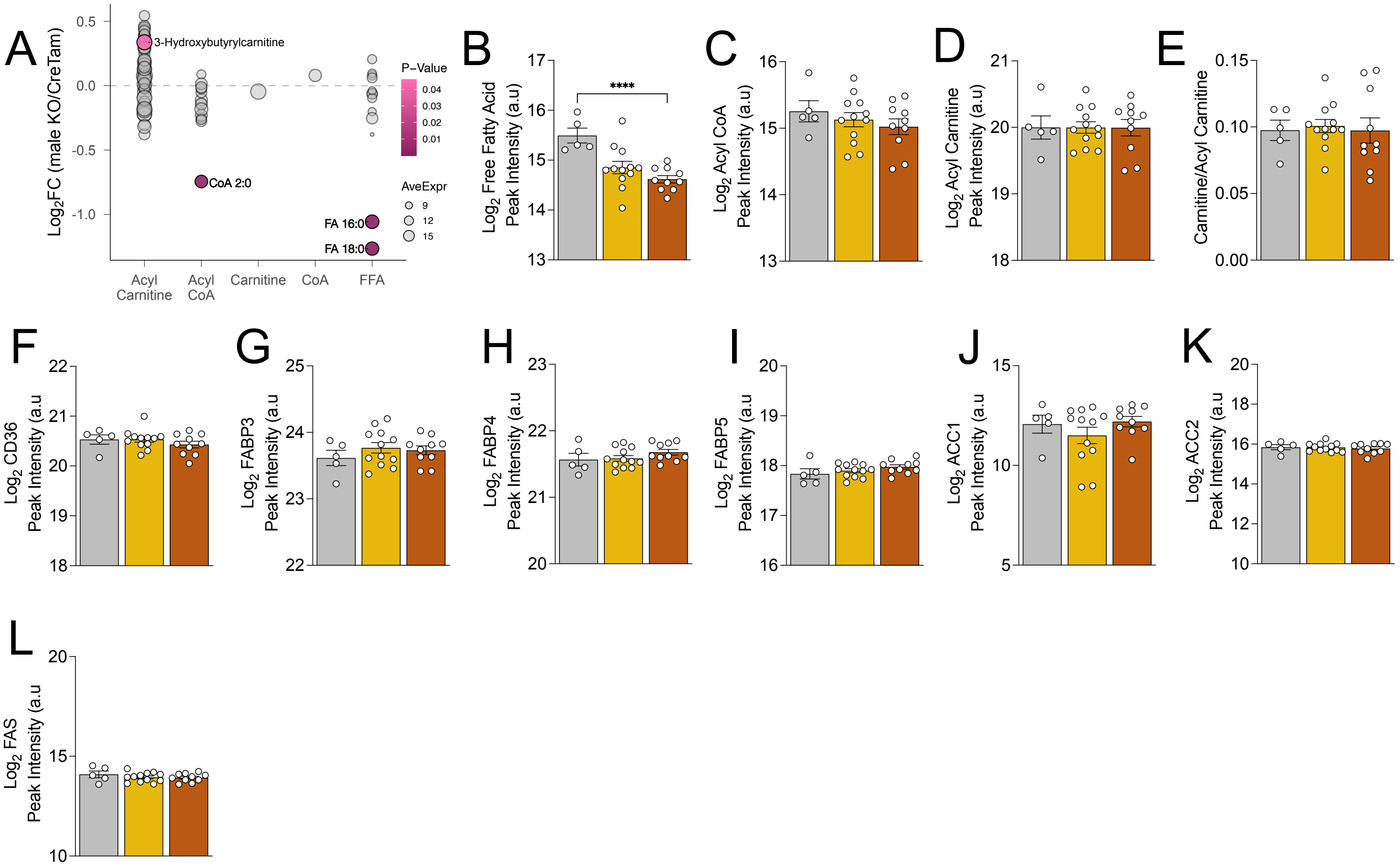


**Fig S7: Effect of PDK1 reduction on male fatty acid metabolism. (A)** Fold change of acyl metabolites in *Pdk1*cKO relative to control in male mice hearts, 2-weeks post-tamoxifen injection. UHPLC-MS/MS based abundance of bulk **(B)** free fatty acids **(C)** acyl CoA **(D)** acyl carnitine **(E)** carnitine to acylcarnitine ratio in female mice**.** Protein abundances of **(F)** CD36; Cluster of Differentiation 36 **(G)** FABP3; Fatty Acid Binding Protein 3 **(H)** FABP4; Fatty Acid Binding Protein 4 **(I)** FABP5; Fatty Acid Binding Protein 5 **(J)** ACC1; Acetyl-CoA Carboxylase 1 **(K)** ACC2; Acetyl-CoA Carboxylase 2 **(L)** FAS; Fatty Acid Synthase. Data are shown as mean ± SEM and were analyzed by either a Limma test (A) or an unpaired two-tailed Welch's t-test (B-E). $^{*}$ $p < 0.05$; $^{**}$ $p < 0.01$; $^{***}$ $p < 0.001$; $^{****}$ $p < 0.0001$.
