## Supplemental Table 1 for "Cardiomyocyte pyruvate dehydrogenase kinase 1 knockout activates Rho-mediated remodeling and decreases fatty acyl availability in female mice"

### Supplementary Tables

**Table S1: Echocardiological parameters.** Data are shown as mean ± SEM. Female CreTam control n = 3, male CreTam control n = 4, female *Pdk1*cKO n = 6, male *Pdk1*cKO n = 3.

| Sex | Female | Female | Female | Female | Female | Female | Male | Male | Male | Male | Male | Male |
|---|---|---|---|---|---|---|---|---|---|---|---|---|
| Genotype | CreTam | CreTam | CreTam | KO | KO | KO | CreTam | CreTam | CreTam | KO | KO | KO |
| Week | Baseline | 2wks | 4wks | Baseline | 2wks | 4wks | Baseline | 2wks | 4wks | Baseline | 2wks | 4wks |
| A' | -16.85 ± 1.36 | -14.17 ± 1.33 | -15.32 ± 3.00 | -14.43 ± 1.29 | -15.84 ± 2.88 | -10.60 ± 1.48 | -14.85 ± 2.51 | -14.55 ± 2.06 | -13.45 ± 1.81 | -14.12 ± 0.98 | -16.14 ± 1.50 | -13.68 ± 0.90 |
| AET | 69.53 ± 3.93 | 66.97 ± 6.14 | 75.82 ± 8.75 | 74.98 ± 1.40 | 69.05 ± 3.43 | 68.75 ± 1.93 | 77.92 ± 5.77 | 69.01 ± 4.50 | 64.33 ± 5.97 | 76.90 ± 5.60 | 67.67 ± 7.02 | 72.04 ± 1.52 |
| E' | -23.90 ± 1.56 | -17.18 ± 3.32 | -19.04 ± 4.13 | -18.77 ± 2.11 | -20.90 ± 3.19 | -15.08 ± 1.61 | -22.74 ± 2.96 | -17.25 ± 2.67 | -18.62 ± 4.94 | -18.65 ± 1.41 | -21.41 ± 1.96 | -20.80 ± 0.87 |
| IVCT | 18.29 ± 1.73 | 18.47 ± 1.39 | 22.51 ± 6.44 | 17.20 ± 2.00 | 16.76 ± 3.88 | 23.72 ± 3.44 | 12.92 ± 1.32 | 18.96 ± 2.34 | 19.90 ± 2.32 | 14.86 ± 1.23 | 14.01 ± 3.05 | 15.19 ± 2.09 |
| IVRT | 23.98 ± 2.91 | 35.80 ± 2.52 | 27.94 ± 1.13 | 19.00 ± 3.03 | 29.61 ± 2.36 | 25.83 ± 2.57 | 13.92 ± 3.54 | 27.19 ± 0.96 | 24.13 ± 0.59 | 17.87 ± 4.57 | 24.60 ± 1.48 | 19.07 ± 7.28 |
| MV A | 465.11 ± 61.7 | 340.77 ± 125 | 386.12 ± 69.1 | 466.71 ± 77.03 | 305.47 ± 46.2 | 302.10 ± 35.9 | 399.51 ± 50.9 | 365.09 ± 32.9 | 321.94 ± 26.73 | 481.17 ± 67.89 | 424.81 ± 34.2 | 387.27 ± 76.17 |
| MV Decel - Acceleration | -58770.88 ± 7 | -55826.37 ± 2 | -45737.72 ± 1 | -48511.96 ± 11555.1 | -36806.52 ± 7 | -38183.12 ± 8 | -33349.83 ± 9 | -40874.47 ± 9 | -29168.95 ± 11083.29 | -45395.35 ± 5377.03 | -38968.48 ± 4 | -35377.91 ± 10002.67 |
| MV Decel - Time | 12.29 ± 1.68 | 13.24 ± 3.65 | 19.03 ± 7.03 | 23.63 ± 7.48 | 18.29 ± 1.78 | 17.08 ± 2.62 | 24.88 ± 6.12 | 21.65 ± 9.41 | 12.78 ± 3.51 | 16.34 ± 1.57 | 19.15 ± 1.05 | 18.70 ± 4.06 |
| MV E | 675.43 ± 45.4 | 476.49 ± 111 | 608.70 ± 72.3 | 722.29 ± 60.24 | 586.33 ± 69.6 | 551.91 ± 54.2 | 657.50 ± 55.3 | 596.24 ± 36.5 | 442.82 ± 122.24 | 707.74 ± 61.87 | 730.91 ± 59.1 | 603.73 ± 134.15 |
| A'/E' | 0.71 ± 0.05 | 0.94 ± 0.20 | 0.81 ± 0.05 | 0.80 ± 0.08 | 0.74 ± 0.03 | 0.70 ± 0.06 | 0.66 ± 0.07 | 0.85 ± 0.04 | 0.80 ± 0.14 | 0.76 ± 0.02 | 0.75 ± 0.01 | 0.67 ± 0.07 |
| E'/A' | 1.43 ± 0.10 | 1.19 ± 0.21 | 1.24 ± 0.08 | 1.31 ± 0.13 | 1.36 ± 0.05 | 1.50 ± 0.15 | 1.57 ± 0.16 | 1.18 ± 0.05 | 1.33 ± 0.21 | 1.32 ± 0.03 | 1.33 ± 0.01 | 1.53 ± 0.15 |
| LV MPI IV | 0.61 ± 0.02 | 0.83 ± 0.08 | 0.68 ± 0.09 | 0.49 ± 0.06 | 0.67 ± 0.04 | 0.72 ± 0.07 | 0.35 ± 0.06 | 0.67 ± 0.02 | 0.70 ± 0.05 | 0.44 ± 0.08 | 0.57 ± 0.04 | 0.47 ± 0.13 |
| MV Area (simplified) | 64.71 ± 7.28 | 74.21 ± 20.45 | 51.59 ± 16.68 | 45.47 ± 10.08 | 43.74 ± 4.68 | 49.35 ± 6.74 | 36.01 ± 7.95 | 51.81 ± 13.63 | 41.29 ± 10.45 | 47.30 ± 4.50 | 39.87 ± 2.15 | 42.54 ± 7.18 |
| MV E/A | 1.51 ± 0.18 | 1.69 ± 0.33 | 1.64 ± 0.19 | 1.72 ± 0.27 | 2.03 ± 0.25 | 1.88 ± 0.18 | 1.72 ± 0.26 | 1.68 ± 0.20 | 1.65 ± 0.11 | 1.49 ± 0.08 | 1.74 ± 0.18 | 1.74 ± 0.06 |
| MV E/E' | -28.76 ± 3.12 | -30.01 ± 6.13 | -34.82 ± 8.27 | -39.57 ± 2.59 | -30.45 ± 4.39 | -38.77 ± 5.44 | -29.52 ± 1.68 | -37.88 ± 7.42 | -33.11 ± 6.93 | -37.90 ± 0.83 | -34.54 ± 3.18 | -30.70 ± 6.15 |
| MV PHT (simplified) | 3.56 ± 0.49 | 3.84 ± 1.06 | 5.52 ± 2.04 | 6.85 ± 2.17 | 5.30 ± 0.52 | 4.95 ± 0.76 | 7.21 ± 1.77 | 6.28 ± 2.73 | 7.38 ± 2.85 | 4.74 ± 0.45 | 5.55 ± 0.31 | 5.54 ± 1.10 |
| AoV VTI- VTI | 149.79 ± 27.7 | 121.23 ± 2.77 | 137.54 ± 12.8 | 86.55 ± 18.42 | 121.07 ± 8.00 | 122.34 ± 13.2 | 128.59 ± 14.1 | 94.24 ± 13.10 | 147.25 ± 40.87 | 120.04 ± 10.85 | 110.22 ± 25.3 | 155.92 ± 25.73 |
| AoV VTI - Mean Vel | 574.80 ± 43.3 | 472.19 ± 72.7 | 442.16 ± 37.6 | 436.22 ± 67.26 | 397.53 ± 21.8 | 455.19 ± 44.9 | 529.27 ± 26.9 | 391.11 ± 59.7 | 531.60 ± 147.81 | 515.93 ± 122.57 | 489.90 ± 96.1 | 707.89 ± 108.27 |
| AoV VTI - Mean Grad | 1.35 ± 0.21 | 0.96 ± 0.31 | 0.79 ± 0.13 | 0.92 ± 0.19 | 0.64 ± 0.07 | 0.87 ± 0.17 | 1.13 ± 0.12 | 0.73 ± 0.16 | 1.39 ± 0.80 | 1.19 ± 0.54 | 1.04 ± 0.41 | 2.11 ± 0.60 |
| AoV VTI - Peak Vel | 1105.25 ± 79 | 897.86 ± 123 | 867.06 ± 90.3 | 843.34 ± 135.81 | 773.51 ± 26.6 | 872.89 ± 77.9 | 1040.53 ± 66 | 746.59 ± 101 | 1096.91 ± 312.43 | 1005.39 ± 191.47 | 913.14 ± 177 | 1340.63 ± 170.54 |
| AoV VTI - Peak Grad | 4.97 ± 0.73 | 3.41 ± 0.99 | 3.07 ± 0.60 | 3.47 ± 0.71 | 2.41 ± 0.17 | 3.17 ± 0.55 | 4.41 ± 0.58 | 2.64 ± 0.42 | 5.98 ± 3.52 | 4.34 ± 1.58 | 3.59 ± 1.40 | 7.42 ± 1.75 |
| AV Peak Vel | 1189.74 ± 12 | 715.34 ± NA | 682.96 ± NA | 805.04 ± 83.94 | 802.34 ± 33.2 | 1085.05 ± NA | 985.90 ± 73.3 | 863.83 ± 56.1 | 771.99 ± 12.22 | 1154.48 ± 198.68 | 909.10 ± 171 | 1423.71 ± NA |
| Desc Ao Vel | -691.03 ± 38. | -500.58 ± 42. | -639.23 ± 19. | -558.40 ± 25.29 | -537.61 ± 30. | -534.85 ± 32. | -768.68 ± 47. | -600.05 ± 61. | -684.52 ± 40.69 | -522.71 ± 76.06 | -625.58 ± 41. | -679.11 ± 21.08 |
| AV Peak Pressure | 5.72 ± 1.18 | 2.05 ± NA | 1.87 ± NA | 2.68 ± 0.59 | 2.59 ± 0.22 | 4.71 ± NA | 3.93 ± 0.60 | 3.01 ± 0.40 | 2.38 ± 0.08 | 5.49 ± 1.83 | 3.54 ± 1.35 | 8.11 ± NA |
| TEI Index | 0.61 ± 0.02 | 0.83 ± 0.08 | 0.68 ± 0.09 | 0.49 ± 0.06 | 0.67 ± 0.04 | 0.72 ± 0.07 | 0.35 ± 0.06 | 0.67 ± 0.02 | 0.70 ± 0.05 | 0.44 ± 0.08 | 0.57 ± 0.04 | 0.48 ± 0.13 |
| IVS;d | 0.86 ± 0.06 | 0.82 ± 0.05 | 0.77 ± 0.10 | 0.71 ± 0.06 | 0.81 ± 0.05 | 0.80 ± 0.02 | 0.73 ± 0.04 | 0.92 ± 0.04 | 0.86 ± 0.07 | 0.79 ± 0.09 | 0.91 ± 0.01 | 0.91 ± 0.06 |
| IVS;s | 1.11 ± 0.06 | 1.08 ± 0.04 | 1.03 ± 0.08 | 1.07 ± 0.04 | 1.01 ± 0.04 | 1.03 ± 0.04 | 0.96 ± 0.03 | 1.16 ± 0.01 | 1.12 ± 0.09 | 1.02 ± 0.08 | 1.23 ± 0.05 | 1.25 ± 0.05 |
| Heart Rate | 416.80 ± 35.5 | 381.52 ± 23.5 | 402.13 ± 31.7 | 471.82 ± 57.51 | 418.19 ± 18.8 | 411.89 ± 20.9 | 409.99 ± 30.1 | 401.62 ± 13.6 | 381.06 ± 19.23 | 390.44 ± 12.71 | 426.14 ± 24.6 | 431.51 ± 23.04 |
| LVID;d | 3.71 ± 0.19 | 3.70 ± 0.14 | 4.07 ± 0.14 | 3.62 ± 0.15 | 3.69 ± 0.12 | 3.60 ± 0.12 | 3.55 ± 0.12 | 3.65 ± 0.05 | 3.86 ± 0.02 | 3.40 ± 0.18 | 3.83 ± 0.14 | 3.67 ± 0.14 |
| LVID;s | 2.43 ± 0.23 | 2.59 ± 0.10 | 2.91 ± 0.14 | 2.25 ± 0.17 | 2.59 ± 0.16 | 2.38 ± 0.10 | 2.21 ± 0.09 | 2.45 ± 0.08 | 2.59 ± 0.10 | 2.22 ± 0.20 | 2.57 ± 0.24 | 2.23 ± 0.12 |
| LVPW;d | 0.70 ± 0.06 | 0.82 ± 0.03 | 0.74 ± 0.04 | 0.69 ± 0.02 | 0.74 ± 0.05 | 0.77 ± 0.03 | 0.67 ± 0.02 | 0.88 ± 0.04 | 0.81 ± 0.07 | 0.75 ± 0.09 | 0.87 ± 0.01 | 0.84 ± 0.05 |
| LVPW;s | 1.06 ± 0.06 | 1.11 ± 0.04 | 1.00 ± 0.08 | 1.06 ± 0.04 | 1.00 ± 0.03 | 1.01 ± 0.03 | 0.99 ± 0.02 | 1.16 ± 0.05 | 1.09 ± 0.11 | 1.00 ± 0.08 | 1.19 ± 0.07 | 1.25 ± 0.03 |
| CO (LV Trace) | 16.44 ± 3.01 | 12.04 ± 1.06 | 15.89 ± 0.96 | 16.14 ± 1.47 | 13.64 ± 1.33 | 14.05 ± 1.33 | 15.26 ± 1.83 | 13.51 ± 0.52 | 14.85 ± 0.56 | 12.34 ± 1.46 | 16.03 ± 1.38 | 16.83 ± 1.04 |
| Diameter;d (LV Trace) | 3.73 ± 0.20 | 3.60 ± 0.11 | 4.06 ± 0.13 | 3.55 ± 0.17 | 3.67 ± 0.13 | 3.61 ± 0.11 | 3.55 ± 0.10 | 3.59 ± 0.04 | 3.82 ± 0.04 | 3.41 ± 0.16 | 3.79 ± 0.12 | 3.63 ± 0.10 |
| Diameter;s (LV Trace) | 2.42 ± 0.21 | 2.53 ± 0.10 | 2.91 ± 0.14 | 2.26 ± 0.18 | 2.58 ± 0.16 | 2.43 ± 0.09 | 2.18 ± 0.10 | 2.41 ± 0.07 | 2.56 ± 0.08 | 2.21 ± 0.16 | 2.56 ± 0.19 | 2.21 ± 0.10 |
| EF | 64.55 ± 4.04 | 57.69 ± 3.44 | 55.70 ± 2.04 | 68.74 ± 3.43 | 56.86 ± 5.28 | 63.38 ± 2.75 | 68.77 ± 1.55 | 62.08 ± 2.58 | 61.75 ± 3.55 | 65.19 ± 4.90 | 61.98 ± 5.35 | 70.44 ± 3.44 |
| EF (LV Trace) | 65.28 ± 2.81 | 57.72 ± 2.94 | 55.01 ± 2.55 | 67.07 ± 3.34 | 57.45 ± 4.69 | 61.69 ± 3.06 | 69.86 ± 2.51 | 62.10 ± 1.76 | 62.12 ± 2.25 | 65.65 ± 4.21 | 61.47 ± 4.20 | 70.32 ± 2.67 |
| FS | 34.82 ± 2.87 | 29.90 ± 2.33 | 28.69 ± 1.26 | 38.13 ± 2.77 | 29.74 ± 3.39 | 33.83 ± 1.95 | 37.73 ± 1.23 | 32.84 ± 1.82 | 32.85 ± 2.53 | 35.07 ± 3.48 | 33.07 ± 3.73 | 39.29 ± 2.80 |
| FS (LV Trace) | 35.39 ± 1.97 | 29.82 ± 1.97 | 28.25 ± 1.60 | 36.77 ± 2.59 | 30.03 ± 3.03 | 32.69 ± 2.12 | 38.70 ± 1.98 | 32.82 ± 1.23 | 33.02 ± 1.61 | 35.43 ± 3.06 | 32.62 ± 2.92 | 39.12 ± 2.16 |
| LV Mass | 101.35 ± 12.6 | 106.93 ± 9.64 | 112.18 ± 10.2 | 83.04 ± 7.25 | 98.60 ± 7.12 | 96.37 ± 3.15 | 80.67 ± 5.64 | 119.74 ± 6.11 | 119.04 ± 13.90 | 85.87 ± 11.88 | 126.20 ± 5.70 | 115.24 ± 3.28 |
| LV Mass (Corrected) | 81.08 ± 10.09 | 85.55 ± 7.71 | 89.74 ± 8.18 | 66.43 ± 5.80 | 78.88 ± 5.70 | 77.09 ± 2.52 | 64.54 ± 4.51 | 95.79 ± 4.89 | 95.23 ± 11.12 | 68.69 ± 9.50 | 100.96 ± 4.56 | 92.19 ± 2.62 |
| LV Vol;d | 59.40 ± 7.77 | 58.41 ± 5.34 | 73.28 ± 5.74 | 55.78 ± 5.81 | 58.09 ± 4.33 | 54.76 ± 4.36 | 53.02 ± 4.07 | 56.26 ± 1.77 | 64.43 ± 0.88 | 47.99 ± 5.85 | 63.22 ± 5.51 | 57.38 ± 5.04 |
| LV Vol;s | 21.80 ± 5.29 | 24.51 ± 2.46 | 32.66 ± 3.93 | 18.01 ± 3.50 | 25.22 ± 3.44 | 20.02 ± 2.02 | 16.58 ± 1.65 | 21.38 ± 1.83 | 24.65 ± 2.31 | 16.96 ± 3.58 | 24.61 ± 5.58 | 16.90 ± 2.23 |
| SV (LV Trace) | 38.60 ± 3.41 | 31.69 ± 2.90 | 39.79 ± 2.45 | 35.23 ± 3.20 | 32.87 ± 3.52 | 34.22 ± 3.19 | 36.90 ± 2.63 | 33.66 ± 0.36 | 39.07 ± 0.89 | 31.59 ± 3.62 | 37.48 ± 1.22 | 39.23 ± 3.13 |
| V;d (LV Trace) | 60.24 ± 8.09 | 55.11 ± 4.16 | 72.65 ± 5.45 | 53.51 ± 6.35 | 57.69 ± 4.42 | 55.31 ± 3.91 | 52.93 ± 3.60 | 54.27 ± 1.43 | 63.07 ± 1.38 | 48.30 ± 5.19 | 61.72 ± 4.88 | 55.75 ± 3.72 |
| V;s (LV Trace) | 21.64 ± 4.71 | 23.42 ± 2.19 | 32.86 ± 3.92 | 18.27 ± 3.63 | 24.81 ± 3.43 | 21.09 ± 1.95 | 16.03 ± 1.96 | 20.61 ± 1.49 | 24.00 ± 1.84 | 16.71 ± 2.98 | 24.23 ± 4.54 | 16.52 ± 1.89 |
| Body weight | 26.30 ± 2.10 | 24.50 ± 1.38 | 28.21 ± 1.83 | 23.25 ± 1.85 | 22.90 ± 0.91 | 25.33 ± 1.49 | 26.90 ± 0.39 | 26.75 ± 0.42 | 29.74 ± 0.39 | 27.57 ± 0.87 | 28.83 ± 1.00 | 32.87 ± 0.98 |
